# Genomic instability within a sympatric complex of South American garlics (Nothoscordum spp., Amaryllidaceae)

**DOI:** 10.64898/2026.06.07.730749

**Authors:** Agostina B. Sassone, Mariela Sader, Thiago Nascimiento, Frank R. Blattner, Liliana M. Giussani

## Abstract

**Background and Aims:** The evolution of reproductive isolation between previously interbreeding populations is a fundamental driver of plant speciation. Within Amaryllidaceae, *Nothoscordum* represents an evolutionarily complex genus, characterized by an unusually high incidence of chromosomal rearrangements. During fieldwork, *Nothoscordum montevidense* and *Nothoscordum bonariense* were found growing in sympatry, along with individuals exhibiting intermediate morphological traits, suggesting a putative hybrid origin. To test this hypothesis, we employed an integrative approach to characterize the morphologically intermediate specimens and the two sympatric populations.

**Materials and methods:** To characterize the putative hybrids we have combined morphological, cytogenetic analyses (chromosome counts, CMA/DAPI banding, and FISH) and flow cytometry-based genome size estimation. Phylogenetic relationships and genomic structure were also investigated through Genotyping-by-Sequencing (GBS), complete chloroplast genome assembly, and comparative repetitive DNA analysis. We also performed species distribution modeling and phenological analyses of the putative parental species.

**Key Results:** Multiple lines of evidence confirm the hybrid origin of the studied plants. Cytogenetic analyses revealed specimens with 2n = 21 (1C ≈ 33 pg = 32.274 Mbp) and 2n = 25 (1C ≈ 37 pg = 36.186 Mbp), accompanied by meiotic irregularities consistent with interspecific hybridization. Chloroplast genome phylogeny identified *N. montevidense* (2n = 16, 1C ≈ 25 pg) as the maternal lineage, while GBS data confirmed *N. bonariense* (2n = 26, 1C ≈ 41 pg) as the paternal contributor and revealed evidence of subsequent backcrossing. Comparative analysis of repetitive DNA showed reduced 35S rDNA diversity in the hybrid, indicative of post-hybridization genomic restructuring. Despite the observed genomic complexity, no clear morphological differentiation was detected among hybrid individuals. Phenological analyses and species distribution models demonstrated broad overlap between parental species.

**Conclusions:** Our findings highlight the role of hybridization in shaping genome architecture in cytogenetically labile plant lineages. Furthermore, our results underscore that morphological similarity can mask profound genomic complexity, reinforcing the value of integrative approaches to understand genera characterized by reticulate evolution and genomic instability.

## INTRODUCTION

Plant speciation occurs when mechanisms of reproductive isolation evolve between previously interbreeding populations (Rieseberg and Willis, 2007). The development of these mechanisms is driven by genomic, structural, and numerical alterations involving either the entire karyotype or individual chromosomes (De Storme and Mason, 2014). In this context, speciation through changes in the complete chromosome complement, such as polyploidy events (autopolyploidy or allopolyploidy), represents the predominant mode of plant speciation, enhancing genetic variation and facilitating adaptation to novel environments (Soltis *et al*., 2015; Cai *et al*., 2019; Lavania, 2020). Hybridization itself is not inherently adaptive; however, its evolutionary impact is more pronounced in groups with frequent hybridization events. According to Ellstrand et al. (1996), such groups are typically characterized by: (a) outcrossing and incomplete reproductive isolation, allowing hybridization between species; (b) developmental and ecological flexibility, enabling hybrids to reach maturity; and (c) perennial growth habits with apomictic or vegetative reproduction, which allow hybrids, even partially sterile ones, to persist and achieve some level of reproductive success. The higher prevalence of hybrid ancestry in plants compared to animals may be attributed more to differences in reproductive isolation mechanisms, developmental flexibility, and opportunities for vegetative reproduction than to the inherently adaptive nature of plant hybrid genomes.

The South American tribe Leucocoryneae (Amaryllidaceae, Allioideae) represents a compelling example for studying the role of karyotype changes as a diversifying evolutionary force (e.g., Souza *et al*., 2016; Sassone *et al*., 2021). Within the tribe, *Nothoscordum* Kunth is the most species-rich genus, with a distribution centered primarily in South America. *Nothoscordum* species are morphologically diverse (e.g., inflorescences ranging from one to 32 flowers), cover a wide ecological range (from the Pampas region to the Tropical Andes, with one species extending into North America), and show remarkable cytogenetic variability. Species in this genus exhibit auto- and allopolyploidy (e.g., Núñez, 1990), a discontinuous series of chromosome numbers based on two basic chromosome numbers, *x* = 4 (4 metacentric) and *x* = 5 (3 metacentric + 2 acrocentric) (Pellicer et al., 2017), and up to six-fold differences in genome size (Sassone et al., 2018). Plants with 2*n* = 8, 10, 16, 18, 19, 20, 24, and 26 chromosomes, varying from acrocentric to metacentric, have been described (Guerra, 2008). The systematics of *Nothoscordum* remain underdeveloped due to several challenges, including a limited number of diagnostic characters for species delimitation, the absence of type specimens for many taxa, and overlapping morphological traits that obscure accurate species identification. These factors complicate the study of biological processes such as speciation and diversification within the genus (Sassone *et al*., 2025). Hybridization and ploidy changes have been proposed as important sources of evolutionary novelty within *Nothoscordum*, fostered by chromosomal rearrangements (e.g. Souza et al., 2019). However, the evolutionary outcomes of hybridization without whole-genome duplication, and the extent to which such hybrids may persist or diversify, remain poorly understood in the genus.

Laboratory crosses conducted among various species of *N.* sect. *Nothoscordum* have demonstrated the existence of low barriers to gene flow within the section (Núñez, 1990). Moreover, Crosa (1974) described the only natural hybrid reported for the genus in Uruguay: *Nothoscordum spathaceum* Parodi (= *N. bonariense* Beauverd) × *N. montevidense* Beauverd (both species belonging to *N.* sect. *Nothoscordum*). Although both species share a distribution across the Pampas region, their ecological preferences typically differ, resulting in predominantly allopatric populations (Crosa, 1974). Nevertheless, the study of natural populations revealed a classic case of intermediate morphology and karyotype consistent with a hybrid origin.

During fieldwork, populations of *Nothoscordum montevidense* and *N. bonariense* were found growing in sympatry, alongside individuals exhibiting intermediate morphological traits. These morphological intermediates were hypothesized to represent natural hybrids. To evaluate the hybrid origin, genomic composition, and evolutionary persistence of these intermediate individuals, we employed an integrative systematic approach combining broad scale phylogenomics with ecological evidence, including species distribution modeling and phenological analyses. The phylogenomic study comprised genome-wide sequencing using Genotyping-by-Sequencing (GBS; Elshire *et al*., 2011) and genome skimming (Kane et al., 2012; Straub *et al*., 2012) of the putative hybrids and their parental species.

## MATERIAL AND METHODS

### Taxon sampling

Populations were collected in October 2012 in the locality of Magdalena (Buenos Aires, Argentina) and maintained in cultivation at the Darwinion Institute of Botany to collect data. Detailed information regarding the voucher specimens is presented in Supplementary Table 1.

### DNA extraction

DNA was extracted from silica-gel dried leaves or fresh leaf tissue (when available) using the DNeasy Plant DNA Extraction Kit (Qiagen, Hilden, Germany) according to the instructions of the manufacturer or with a modified CTAB procedure (Doyle and Doyle, 1987). DNA concentration and quality were afterwards checked on 0.8% agarose gels. Afterwards, all DNA samples were quantified using the Qubit dsDNA BR Assay kit (Life Technologies) on a Qubit v2.0 (Life Technologies).

### Whole Genome Shotgun Sequencing

We obtained the complete plastome of South American Allioideae using genome skimming by low-coverage whole-genome shotgun sequencing. Genomic DNA were generated using the Illumina Nextera® DNA Flex Library Preparation Kit (Illumina, San Diego, California, USA). Initially, 30 ng of the plastid DNA was fragmented, cleaned, and amplified, and libraries were prepared as per the manufacturer’s protocol. Barcodes were added, and 500–600-bp fragments were size-selected after quality assessment using an Agilent Bioanalyzer (Agilent Technologies, Santa Clara, California, USA) and qPCR. Multiplexing using ∼250 bp paired-end chemistry was performed in one lane on Illumina NovaSeq 6000.

### Chloroplast Genome Assembly

Genomic raw sequencing reads were quality-checked and assessed for over-represented sequences using FastQC (Andrews, 2010). Error correction and removal of clonal reads were carried out using BBTools (sourceforge.net/projects/bbmap/). The filtered reads were then de novo assembled into contigs using GetOrganelle v.1.7.5 (Jin *et al*., 2020), which was used to generate draft assemblies from 2×250 bp Illumina paired-end reads. Initial annotation of the chloroplast genomes was conducted using GeSeq v2.03, including the annotation of inverted repeats (IRs), the *rps12* interspersed gene, protein-coding sequences, transfer RNAs (tRNAs), and ribosomal RNAs (rRNAs), applying identity thresholds of 55% for protein annotation and 90% for DNA and RNA sequences. tRNAscan-SE v2.0.7 and Chloë v0.1.0 (https://github.com/ian-small/chloe) were additionally incorporated into GeSeq as supplementary annotation tools. All annotations were subsequently manually refined using Geneious Prime v2023.2.1. Multiple sequence alignments were performed using MAFFT v7.245 (Katoh and Standley, 2013) with default settings. Phylogenetic relationships were inferred by a maximum-likelihood analysis using the GTR+Γ substitution model using the complete plastid data with only one IR, including 23 taxa (ingroup + other Allioideae specimens acting as outgroups, Table S1). The tree search was conducted through an estimation of the proportion of invariable sites, and a total of 100 nonparametric bootstrap replicates were performed in RAxML *v*. 8.2.9 (Stamatakis, 2014). Results were summarized using a 50% majority-rule consensus tree.

### Genotyping-by-Sequencing

For library preparation, 20 ng of genomic DNA was used and cut with two restriction enzymes, *Pst*I--HF (NEB) and *Msp*I (NEB). Individual barcoding and single-end sequencing were performed on the Illumina NovaSeq 6000 following Ala *et al*. (2025). GBS library construction and sequencing were performed at the Leibniz Institute of Plant Genetics and Crop Plant Research (IPK), Germany. Quality assessment of all raw sequence samples was performed using FastQC (Andrews, 2010).

### GBS Data assembly, data exploration

GBS loci were assembled using the ipyrad v. 0.9.97 pipeline (Eaton and Overcast, 2020). The running parameters were set according to default recommendations (available in the ipyrad documentation) and explored using settings after Eaton *et al*. (2017) and Gargiulo *et al*. (2021). Briefly, assemblies were generated employing clustering thresholds c = 0.85 and a minimum number of between 4 and 15 samples per locus, depending on the desired subsequent analyses. Statistical base calling was conducted with a maximum of five *Ns* in consensus sequences. To explore single-nucleotide polymorphism (SNP) data, a PCA was performed using the ipyrad application programming interface (API) (Eaton and Overcast, 2020). The potential effect of linkage between SNPs was reduced by subsampling one SNP per locus.

### Phylogenomic analyses

Phylogenetic relationships were inferred by a maximum-likelihood analysis using the GTR+Γ substitution model and a data set with 15 ingroup samples (*N. bonariense* + *N. montevidense* + putative hybrids or morphological intermediates) plus 14 specimens of other *Nothoscordum spp*. acting as outgroups (data set 1, Table 1). The tree search was conducted through an estimation of the proportion of invariable sites, and 1000 nonparametric bootstrap replicates were performed in RAxML *v*. 8.2.9 (Stamatakis, 2014). Results were summarized using a 50% majority-rule consensus tree.

**Table 1.** Next-generation sequencing characteristics of the four assembled Genotyping-by-sequencing data sets under ipyrad version 0.9.97.

| Dataset | Samples | No. of loci | Concatenate<br>d length (bp) | % Missing<br>data | No. SNPs | Minimum samples<br>per locus |
| --- | --- | --- | --- | --- | --- | --- |
| 1 | 29 | 18167 | 1726906 | 71.68 | 95725 | 4 |
| 2 | 15 | 1947 | 185322 | 20.44 | 5219 | 10 |
| 3 | 15 | 220 | 21282 | 0.61 | 476 | 15 |

To provide an alternative representation of evolutionary relationships, we constructed a Neighbor-Net phylogenetic network using a distance-based approach. The network was generated from uncorrected P-distances, calculated as the proportion of nucleotide sites differing between pairs of sequences without applying a substitution model (Nei and Kumar, 2000), using SplitsTree5 v. 5.1.4-beta (Huson and Bryant, 2006). The underlying distance matrix was derived using the Neighbor-Joining algorithm of Saitou and Nei, as implemented by Bryant and Moulton (2004). Neighbor-Net networks can simultaneously represent multiple conflicting tree topologies, making them particularly informative for detecting hybridization or reticulation events (Bryant *et al*., 2007).

### Genetic structure

The Bayesian clustering method STRUCTURE v. 2.2.4 (Pritchard *et al*. 2000) was used to determine the number of distinct genetic clusters (K) with a burn-in period of 500,000 repetitions followed by 2,000,000 repetitions, as implemented in the ipyrad API. Fifteen replicate analyses (data sets 2 and 3) were performed for each data set with values of K = 1–10.

### Morphological assessment

Based on a previous morphological multivariate analysis, Sassone *et al*. (2013) used 51 characters to test phenetic similarity within the Leucocoryneae. In this work, morphological exploration was limited to 24 characters (from which 7 were qualitative traits) that present variation within *Nothoscordum* and related species. We measured three specimens collected in the field per species and the putative hybrids (intermediate morphological specimens) as well, as listed in Table S2.

### Phenology

Information on flowering times was retrieved from labels of 129 herbarium specimens stored at SI (repository link) and plotted using packages “car” (Fox, 2019) and “circular” (Agostinelli, 2025) implemented in Studio (Posit team, 2025).

### Cytogenetic Techniques

The mitotic chromosome preparations were obtained from root meristems pretreated with 0.2% colchicine at 10 °C for 24 h and fixed in ethanol/acetic acid (3:1, v:v). The tissues were digested with 2% (w/v) cellulase (Onozuka)/20% (v/v) pectinase (Sigma) solution at 37 °C for 60 min and squashed in 45% acetic acid. Preparations were frozen in liquid nitrogen to remove the coverslip. To prepare meiotic chromosomes, inflorescences were fixed as described above for roots. Meiotic spreads were prepared by a typical squash method.

For CMA/DAPI staining, the slides were aged for 3 days, stained with CMA 0.1 mg/mL for 30 min, and mounted in glycerol/McIlvaine buffer pH 7.0 (1:1) containing 2.5 mM MgCl2 and 1 μg/mL DAPI. The slides were then maintained in the dark at room temperature for 3 days before analysis. Fluorescence *in situ* hybridization (FISH) was carried out following protocols by Schwarzacher and Heslop-Harrison (2000). To reach stringency above 76%, the hybridization mix had 2x SSC, 50% v/v formamide, 20% v/v dextran sulfate, 0.1% v/v SDS, and 4–6 ng/μL probes. Post-hybridization washes consisted of 2x SSC, 0.1x SSC, and 2× SSC, for 10 min at 42 °C each. The 5S and 35S rDNA probes were designed by Waminal *et al*. (2018) and labeled in Macrogen, Inc. (Seoul, Republic of Korea). Slides were analyzed and photographed using the Leica DMLB epifluorescence microscope, and images were captured with a Cohu CCD video camera using Leica QFISH software and adjusted uniformly for brightness and contrast in Adobe Photoshop CS5 v.12.0.

### Flow cytometry

The DNA content of the specimens was estimated using fresh leaves on a CyFlow Space (Sysmex Partec, Germany) flow cytometer against *Ipheion uniflorum* (Graham) Raf (19.3 pg 2C) or *Vicia faba* L. as internal standard (26.5 pg 2C), respectively. We used the “CyStain PI Absolute P kit” (Sysmex Partec, Görlitz, Germany) nuclei isolation buffer following the manufacturer’s instructions. All cytometric parameters (chromatogram peaks, mean values, and coefficient of variation) were calculated using FloMax® software (Sysmex Partec GmbH, Münster, Germany).

### High-throughput sequencing, data processing, and repetitive DNA identification

All sequences were quality-filtered using a cut-off value of Q20 and retaining 90% of bases equal or above this value using the FASTQ/A short-reads pre-processing tools (Gordon; 2010; http://hannonlab.cshl.edu/fastx_toolkit/) as implemented in RepeatExplorer2. The similarity-based clustering analysis was performed using the RepeatExplorer2 pipeline provided by the ELIXIR-CZ project, part of the international ELIXIR infrastructure (https://www.repeatexplorer-elixir.cerit-sc.cz) (Novák *et al*. 2013, Neumann *et al*. 2019), with no read sampling selection during the automatic sampling step. A comparative analysis involving the simultaneous clustering of reads from all accessions was performed following the protocol of Novák and Neumann (2020) with default settings. Sequencing reads were jointly clustered to identify shared repetitive elements and assess differences in their relative abundances. Reads from each accession were labeled with unique prefixes, concatenated, and processed through the same pipeline using the comparative analysis option (Table S3). From this analysis, the distribution of the most prevalent comparative repeat clusters was graphically represented, excluding clusters originating from plastid-derived sequences. The final genome coverage used for clustering after automatic sampling was calculated as follows: coverage = (r×l)/g, where r corresponds to the number of analyzed reads after clustering, l to read length, and g to haploid genome size in bp.

The TEs were automatically annotated using the green plants (Viridiplantae 4.0) database in RepeatExplorer2. This classification system reflects the elements’ phylogenetic relationships and distinct sequence and structural features (Neumann *et al*., 2019). A database of retrotransposon protein domains (REXdb) is implemented in the RepeatExplorer web server. Repeat composition was calculated excluding clusters of organelle DNA, probably representing extranuclear DNA from chloroplasts and mitochondria. Although comparative repeat analysis provides a genome-wide overview of repetitive DNA, in this study, it was mainly explored to assess patterns of ribosomal DNA (5S and 35S rDNA) abundance and divergence among accessions, following previous cytogenomic approaches (e.g. Garcia *et al*., 2020).

### Distribution map and species distribution modeling

Distribution maps were constructed from georeferenced sites obtained from field collections as well as herbarium specimens stored at the herbarium of the Darwinion Institute (SI), and available at the repository (link). This resulted in a total of 83 occurrences for *N. montevidense* and 39 for *N. bonariense* plus 2 localities (one from our study and one reported by Crosa, 1974) with intermediate morphology (putative hybrids). The distribution map was assembled from the georeferences using the package “raster” (Hijmans *al*., 2015). To test Crosa’s (1974) hypothesis of the species not sharing ecological preferences, we retrieved data for the 19 bioclimatic variables from WorldClim 2.1 (Fick and Hijmans, 2017) with 2.5 min (∼5 km2) spatial resolution. The species distribution models were constructed using MaxEnt v.3.4.3 (Phillips et al., 2006) in the R environment (R Core Team, 2017) and ‘raster’ (Hijmans *et al*., 2015) and ‘rJava’ (Urbanek, 2015). To avoid cross-correlation within climatic variables for the complete extent of the distribution area of the complex, a multicollinearity test was conducted between all pairs of variables using Pearson’s correlation coefficient. Correlations were analyzed in ‘raster’ and plotted as pairs using ‘corrplot’ (Wei and Simko, 2024, Fig. S1 and S2). By selecting variables with low correlation coefficients (< 0.5), multicollinearity was minimized. A raster grid stack of the variables for the area of interest was generated and the relevant data at each collection point were extracted using the R package ‘dismo’ (Hijmans *et al*., 2024). The following settings for the model were used: the replicates with Bootstrap, response curves, jackknife tests, logistic output format random seed, random test percentage of 10% to compute the area under the curve (AUC), and to determine the threshold value a 10-percentile training presence was selected. Five of the 19 available bioclimatic variables were used for the final prediction models.

All cited packages were implemented in the R environment and used in RStudio. Computational infrastructure and support were provided by the Central Information Technology at IPK Leibniz Institute.

## RESULTS

### Morphological description

Examination of the the putative hybrids collected in the field revealed intermediate traits between *N.bonariense* and *N. montevidense*. Particularly in flower coloration, which ranged from cream to pale yellow (Fig. 1B), compared to the distinct white and yellow flowers from *N. bonariense* and *N. montevidense,* respectively (Fig. 2A). Plant height of those with intermediate morphology also fell between that of the parental, ranging from 14 to 21 cm, as did the number of flowers per inflorescence (Table S2). All the putative hybrids exhibited rhizomes, a characteristic shared with *N. bonariense*. The intermediate specimens displayed a receptive stigma, along with the presence of both pollen and ovules (Fig. 2, Table S2); however, although these individuals produced fruits across different seasons, these fruits consistently lacked seeds. Additionally, like both parental species, the intermediate specimens also exhibited ligules on their leaves (Guaglianone 1972), which is not a common trait of the subfamily Allioideae but seems frequent in a clade within *Nothoscordum* section *Nothoscordum* (ongoing studies) where *N. montevidense* and *N. bonariense* are included.

**Fig. 1.**
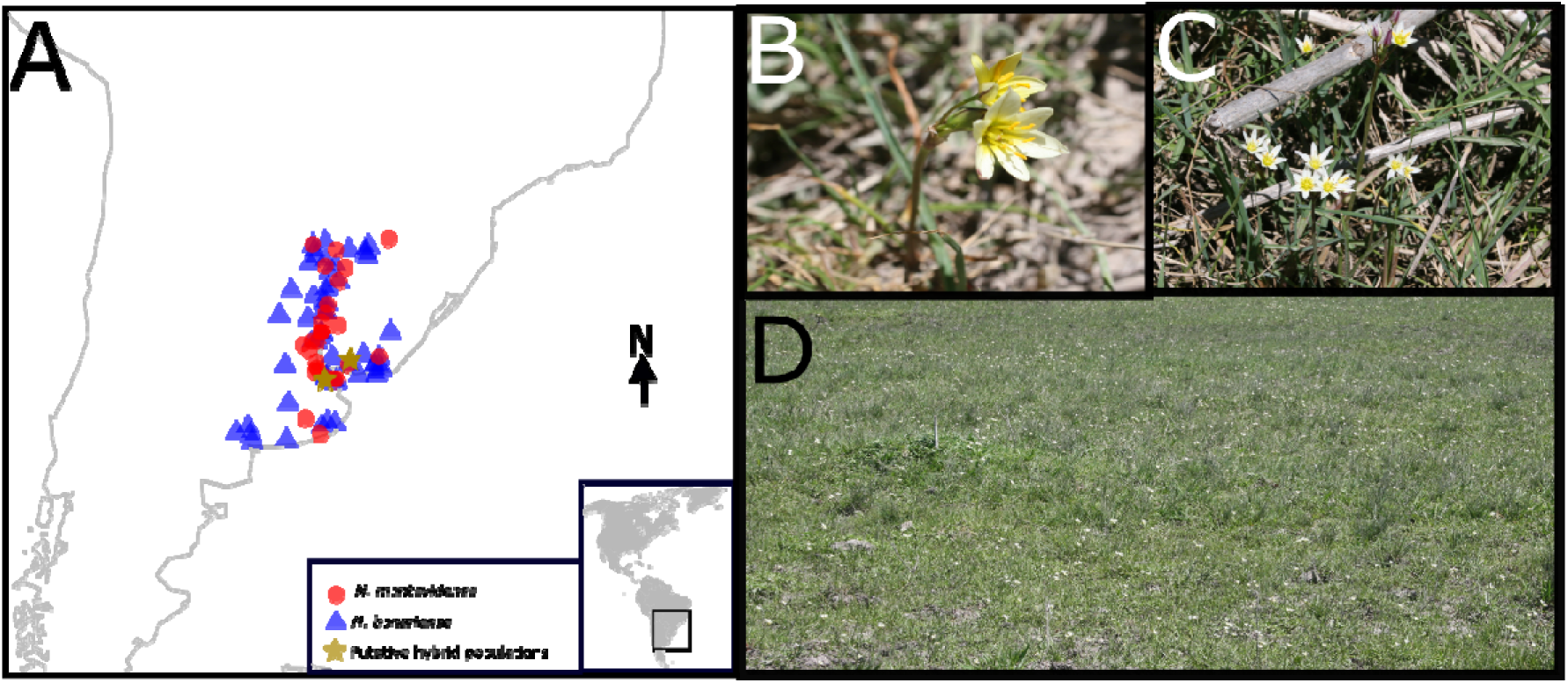
A. Geographical distribution of the focused species. Stars indicate the two reported populations of possible hybridization. B.-C Natural population of putative hybrids of *Nothoscordum* D. Studied population in Magdalena locality, Buenos Aires province (Argentina). Photo credits: B-D: Foto taken by L. M. Giussani- O. Morrone.

**Fig. 2.**
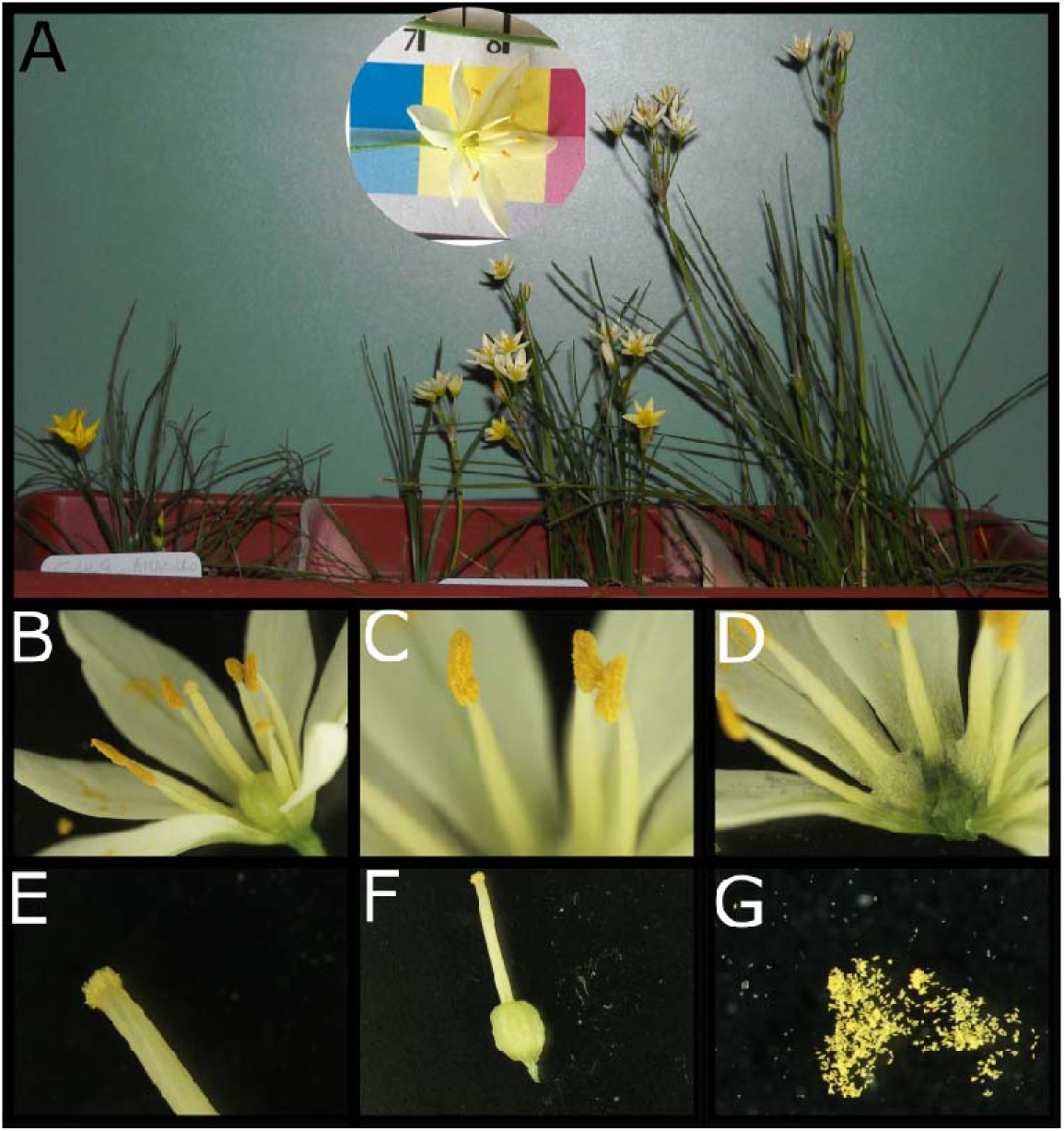
Morphological features. A. Specimens in cultivation from the described collection site, representatives of the 3 populations found, from left to right: *N. montevidense* (*Giussani, L. 449*), the hybrid *Nothoscordum montevidense* x *N. bonariense* (*Giussani, L. 448*), and *N. bonariense* (*Giussani, L. 450*). B-G: Putative hybrid. B. Open flower C. Staminal filaments D. Open flower showing the presence of nectar, E.Receptive style F. Gynoecium G. Pollen.

### Phenology

*Nothoscordum bonariense* and *N. montevidense* exhibit two flowering periods annually; a higher flowering frequency during spring and a second flowering during autumn. Both species showed the main flowering frequency between October and December (Fig. 3; Flowering frequency was plotted as a circular histogram). Flower longevity in both species extends to ca. 4-5 days, and *Nothoscordum bonariense* is a species capable of autogamy.

**Fig. 3.**
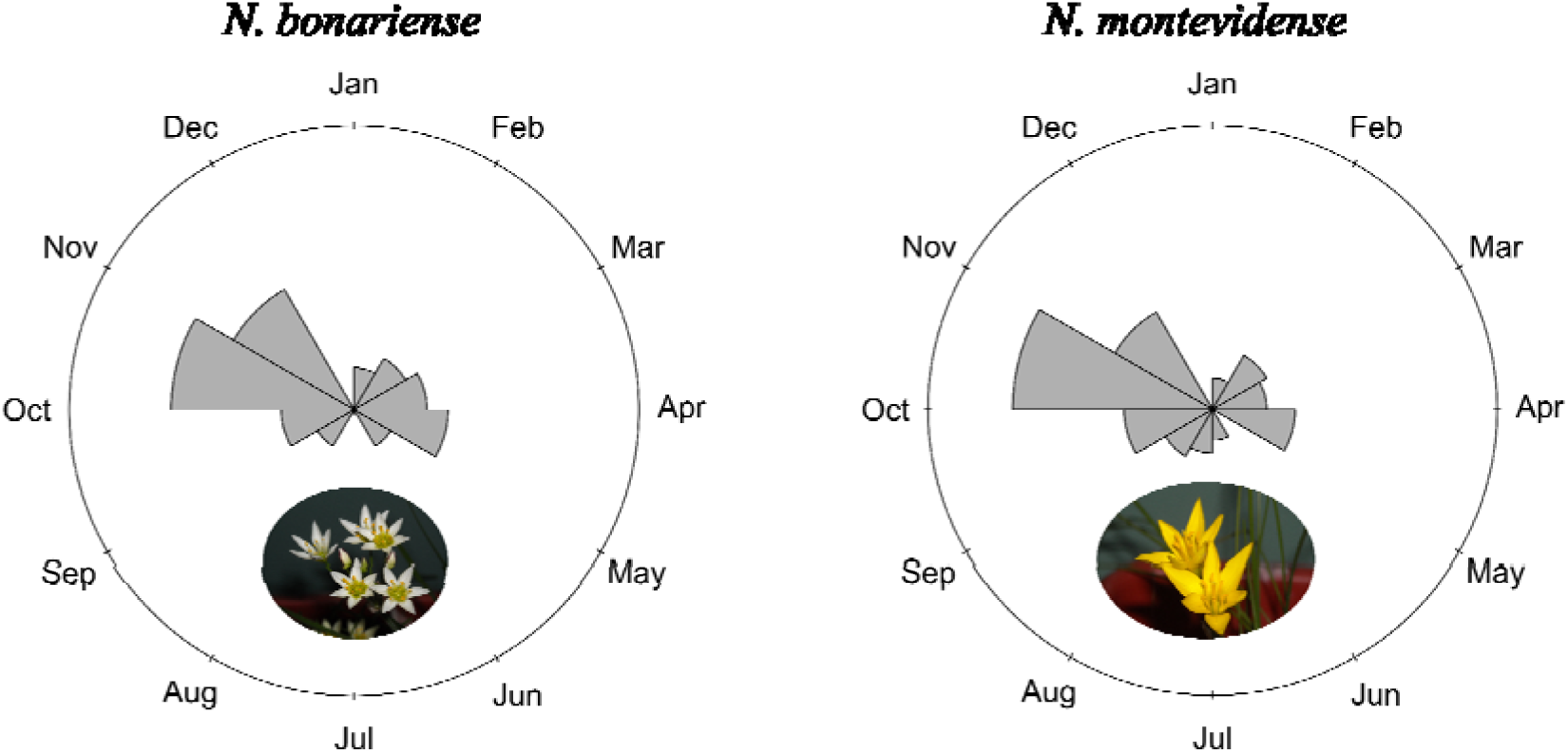
Circular histograms of flowering phenology for *Nothoscordum bonariense* and *N. montevidense*.

### Species distribution modelling

The species distribution models mostly reflect the known distribution of each species. The average AUC test values of the SDM models were 0.936 (*N. montevidense*) and 0.970 (*N. bonariensis*) for 10 repetitions. Although both potential distributions overlap, *N. montevidense* exhibits a higher habitat suitability in the northeastern area (Fig. 4A) than *N. bonariense* (Fig. 4B). The models showed that the bioclimatic variables contributing to the species models are shared for both species. bio14 (Precipitation of Driest Month), bio 4 (Temperature Seasonality) and bio 19 (Precipitation of Coldest Quarter) contributed the most to the potential distribution of both species (Table S4 and S5).

**Fig. 4.**
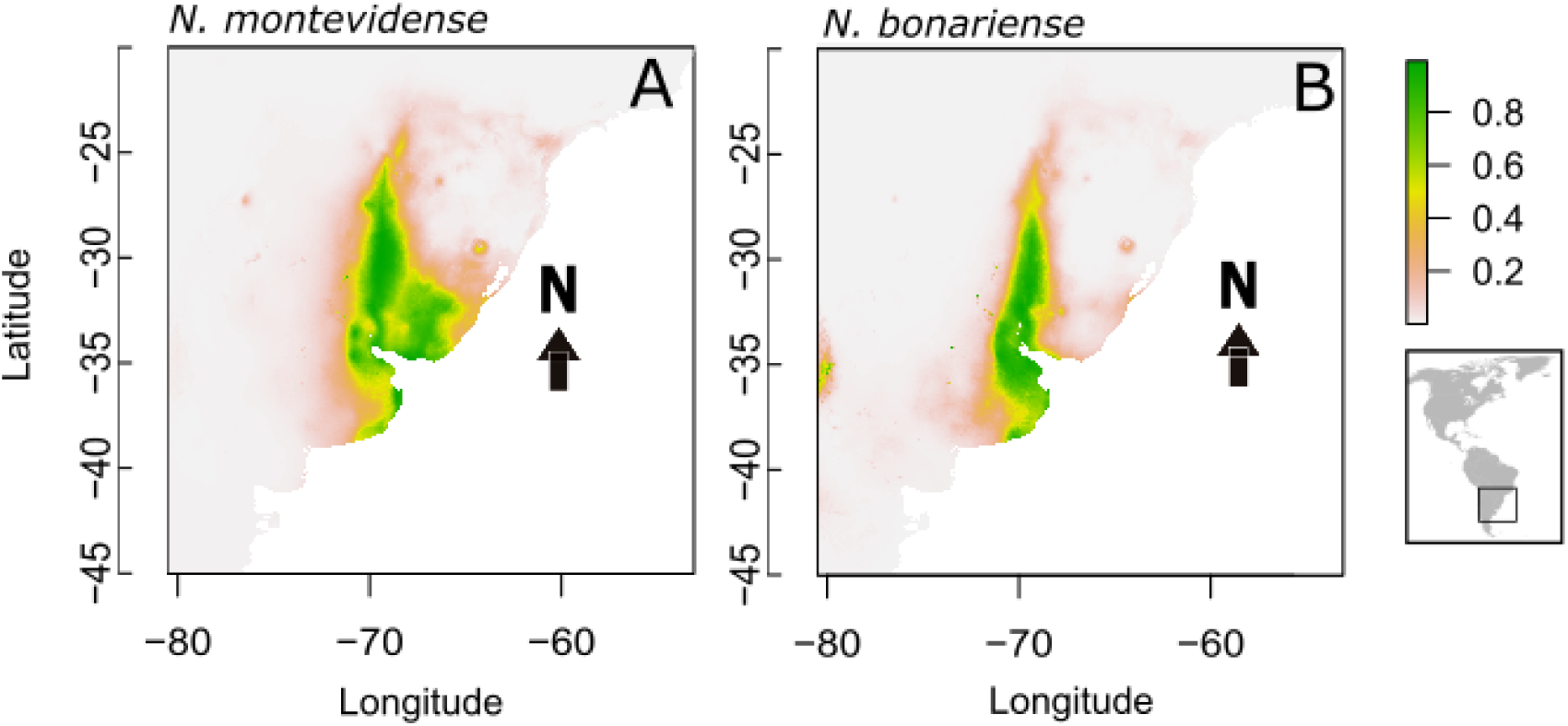
Species distribution modeling for *Nothoscordum montevidense* (A) and *N. bonariense* (B) based on MaxEnt v.3.4.3 analysis as implemented in the R environment. The maps show the predicted suitable habitats for both species. White to green areas indicate non-suitable to suitable habitat.

### Cytogenetic analyses

To determine the distribution of heterochromatin (GC-rich regions), double staining with the fluorochromes CMA and DAPI was performed in the putative hybrids. Chromosome counts revealed individuals with 2*n* = 21 chromosomes, which consistently exhibited two pairs of terminal CMA-positive bands located on the short arm of the second metacentric chromosome pair and on the acrocentric chromosome pair. Less frequently, individuals with 2*n* = 25 chromosomes were also detected (Fig. 5). No complex heterochromatin patterns or DAPI-positive bands were observed in any of the analyzed karyotypes.

**Fig. 5.**
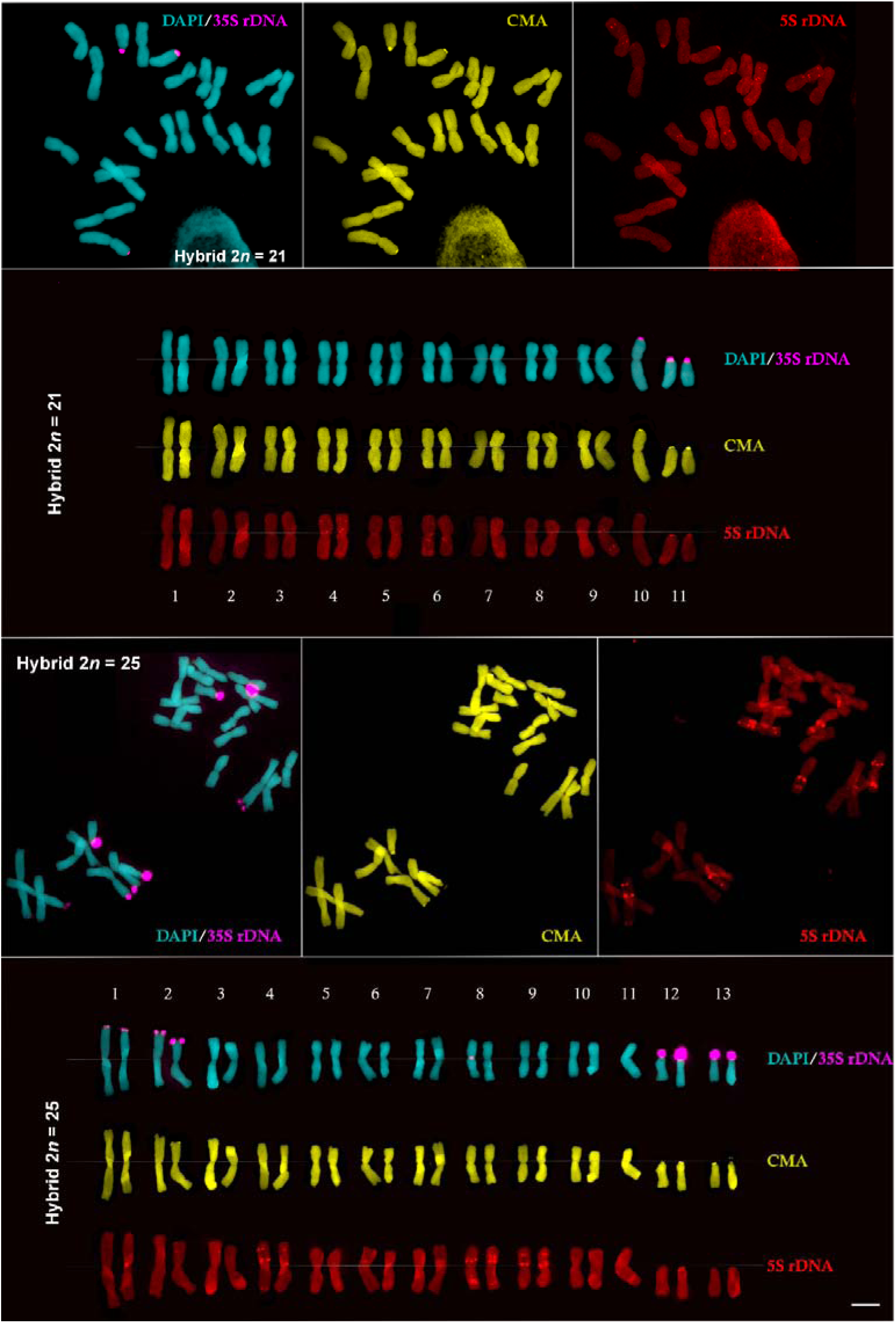
Mitotic metaphase chromosome spread and corresponding karyotype of the hybrid *Nothoscordum montevidense* × *N. bonariense* (Giussani, L. 448), with 2*n* = 21 and 2*n* = 25 chromosomes. Chromosomes were counterstained with DAPI (blue), GC-rich heterochromatin was detected by CMA staining (yellow), and ribosomal DNA sites were localized by FISH using 5S rDNA (red) and 35S rDNA (pink) probes. The lower panels show the corresponding karyotype. Bar = 10 μm.

FISH analysis showed that karyotypes with 2*n* = 21 possessed two pairs of terminal 35S rDNA signals co-localized with CMA-positive bands. One pair was located at the end of the short arm of the second metacentric pair, whereas the other was detected on the acrocentric pair. In addition, seven interstitial pairs of 5S rDNA sites were observed on metacentric chromosomes, together with one pair on the acrocentric chromosome (Fig. 5).

FISH analysis of individuals with 2*n* = 25 revealed four pairs of terminal 35S rDNA signals co-localized with CMA-positive bands. One signal pair was located at the end of the short arm of the second metacentric pair, whereas the other pair, together with the unpaired signal, was associated with the acrocentric chromosomes. The distribution of 5S rDNA sites was identical to that observed in individuals with 2*n* = 21, with seven interstitial pairs on metacentric chromosomes (Fig. 5).

Meiotic analyses revealed irregular meiosis, characterized by lagging chromosomes during anaphase and frequent micronucleus formation (Fig. S3).

### Genome size

The results of the estimation of the DNA content are summarized in Table 2. *Nothoscordum montevidense* exhibited a genome size of ca. 1C ≈ 25 pg, whereas *N. bonariense* ca. 1C ≈ 41 pg. Among the putative hybrids, two specimens measured 1C ≈ 32.7–33 pg, whereas the remaining specimens ranged from 1C ≈ 36.5 to 38.5 pg.

**Table 2.**
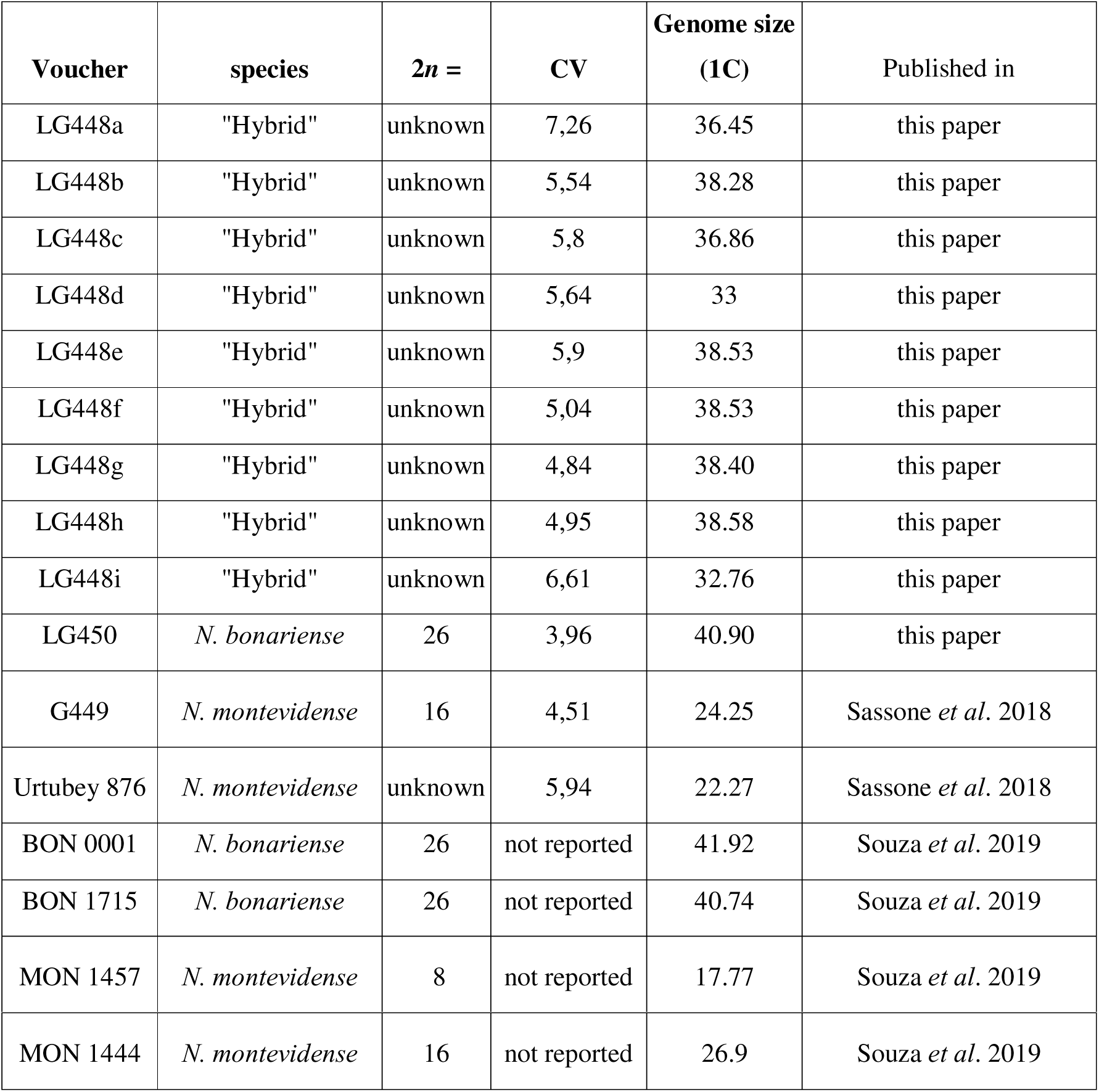
Cytological analyses and DNA content of *Nothoscordum montevidense*, *N. bonariense* and the putative hybrids. CV: Coefficient of Variation en %.

### Repeats

To further characterize the rDNA fraction across species, we performed both individual and comparative analyses of repetitive sequences (Fig. 5; Table S3). The proportion of 5S rDNA was similar across both species and the putative hybrid, representing approximately 0.02% of the genome. In contrast, 35S rDNA accounted for 0.33% in the hybrid, 0.56% in *N. montevidense*, and 0.61% in *N. bonariense*.

Individual graph-based analyses of 35S rDNA in the parental species revealed three clusters in *N. bonariense* and two clusters in *N. montevidense*. The comparative analysis, incorporating reads from both parental genomes and the hybrid, identified two distinct clusters, each containing reads from all three genomes; however, hybrid-derived reads were consistently represented at lower proportions (Cluster 160: 0.16%; Cluster 177: 0.13%). In contrast, the individual analysis of the hybrid recovered only two clusters, accounting for 0.14% and 0.082% of total read abundance. This pattern suggests reduced rDNA cluster diversity in the hybrid relative to *N. bonariense*, consistent with the differential retention or loss of parental rDNA variants following hybridization.

The individual graph-based analysis of 5S rDNA in the hybrid yielded a clustering pattern identical to that recovered in the comparative analysis incorporating reads from both parental species and the hybrid. In both cases, we observed extensive read intermixing and shared cluster topology, supporting the contribution of both parental genomes to the hybrid and confirming its hybrid origin (Fig. 6).

**Fig. 6.**
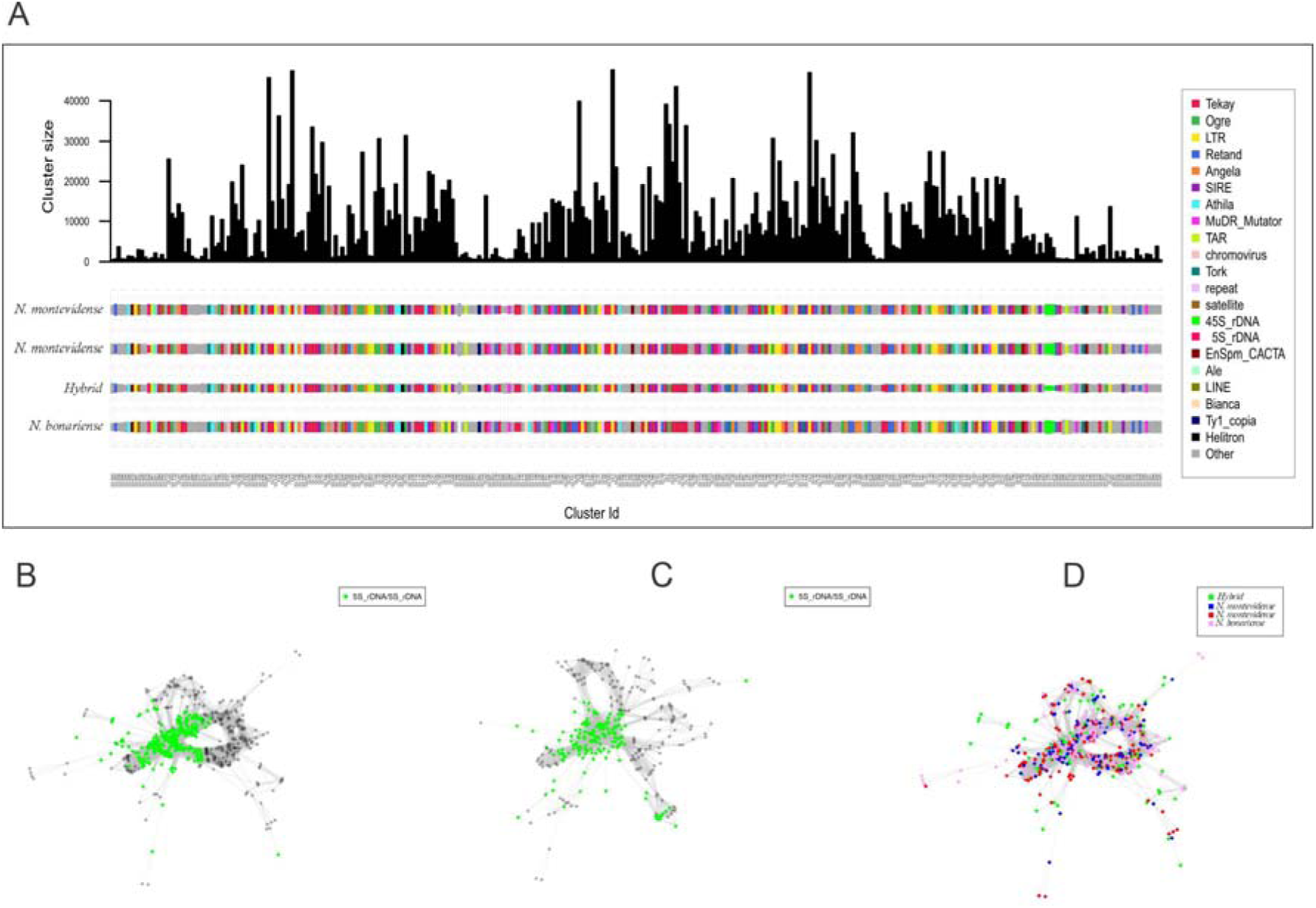
A. Comparative genomic proportions (%) of repetitive sequence families across analyzed *Nothoscordum* populations obtained through the comparative clustering analysis in RepeatExplorer2. B. Graph layout from the comparative analysis using the parental and hybrid reads showing the 5S rDNA (green) interconnected with grey nodes representing an intergenic spacer. C. Graph layout from the individual analysis using the hybrid reads showing the 5S rDNA (green) interconnected with grey nodes representing an intergenic spacer. D. Graph layout from the comparative analysis using the parental and putative hybrid reads showing the 5S rDNA clusters, with colored nodes representing reads from each accession.

### Phylogenetic backbone

The aligned matrix of the plastome data resulted in 160,043 bp length based on 23 taxa, including outgroups; of these 125,424 (79.7%) were uninformative sites. The phylogeny showed that two specimens of *N. montevidense* from our studied population shared an identical chloroplast with the putative hybrid, suggesting this species as the maternal lineage (Fig. 7). As regards *N. bonariense*, the specimen from the Magdalena population (sympatric with the putative hybrid) clustered together with specimens from different populations of the Pampean region: *N. bivalve* (previously proposed to be a progenitor of this species), *N. montevidense* from Uruguay, and *N. minarum*. If we look at the similarity of the plastid sequences, the hybrid and *N. montevidense* shared the same chloroplast, and 98.25% of similarity with *N. bonariense*.

**Fig. 7.**
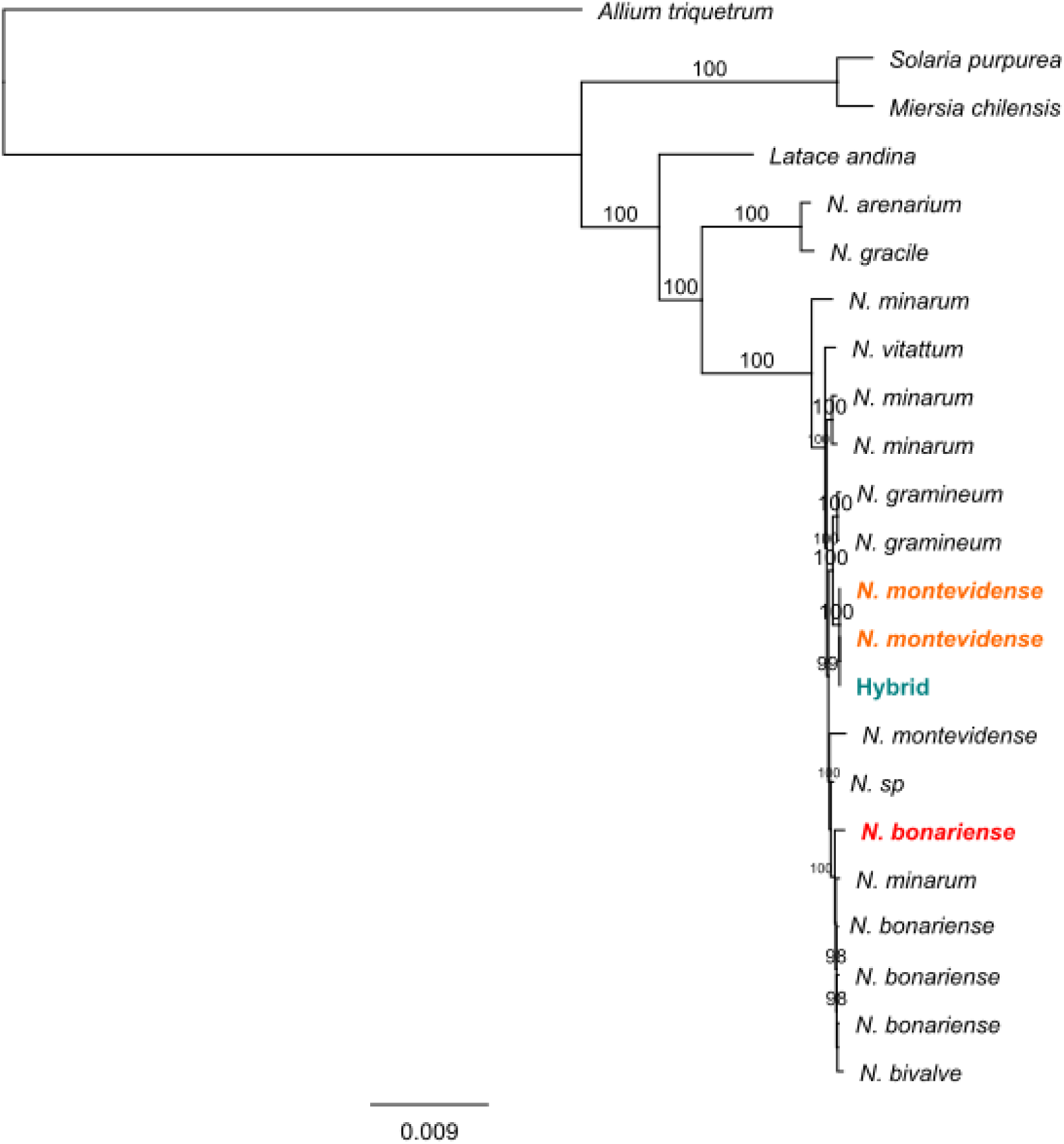
Plastome phylogeny of 23 Allioideae taxa inferred using maximum likelihood implemented in RaxML *v*. 8.2.9. Bootstrap values below 98 are not shown.

### Genomic variability based on GBS data

The first two components of the PCA of a subsampling of one SNP per locus (data set 3, Table 1) explained 54.3% of the observed variance, showing that the parental species as well as the putative hybrids are well-differentiated (Fig. 8A). The intermediate specimens were differentiated from *N. bonariense* and *N. montevidense*, forming three distinct groups that could be distinguished among themselves. In the analysis of the SNP-based population structure, Evanno’s ΔK method suggested K = 2 and the next best fit at K = 3 (Fig. S4), identifying compositional differences among species and with the putative hybrid. Two specimens of the putative hybrid (1C ≈ 33 pg) are genomically more similar to *N. montevidense* than the rest of the specimens (Fig. 8D) and another one more similar to *N. bonariense* than the rest. Uncorrected P- distance-based split networks of the SNP matrix recovered species groups and displayed a reticulated pattern (Fig. 8B).

**Fig. 8.**
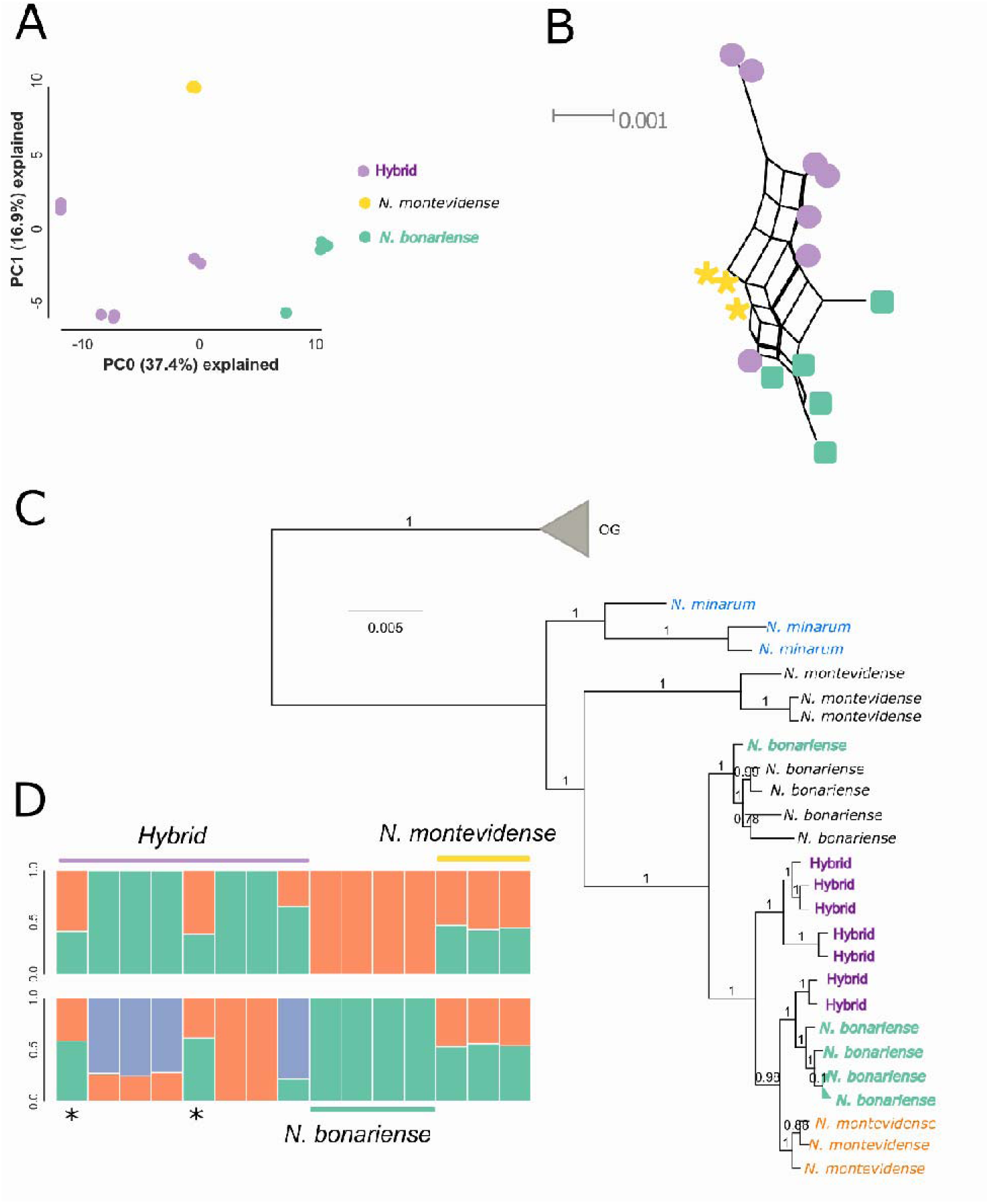
Analysis of Genotype-by-sequencing of specimens of *Nothoscordum* species from the focused populations (15 specimens): A. Principal Component analysis, first and second principal components of dataset 4, based on 1,003/2,947 unlinked SNPs. B. Phylogenetic network analysis calculated using the Neighbor-Net algorithm. Each symbol represents an individual. C. Maximum-likelihood inference for 29 samples, representing sampled populations and outgroups. Bootstrap values are indicated above branches. D. Structure analyses based on based on 1,003/2,947 unlinked SNPs (datset 4). Individuals are represented by a vertical bar that is divided by colored segments representing the likelihood of membership in each cluster, Evanno’s ΔK method suggested K = 2 and the next best fit at K = 3, asterisks (*) mark intermediate specimens genetically closer to *N. montevidense*.

The phylogenetic inference showed that all accessions from Magdalena locality grouped together in a clade. Only one out of 5 studied accessions of *N. bonariense* from Magdalena locality grouped apart with other group of specimens of *N. bonariense* from different localities (Fig. 8C).

## DISCUSSION

Hybridization plays a significant role as an evolutionary force, contributing to plant speciation and diversity. It can occur at both homoploid and polyploid levels, with polyploidy being particularly impactful in generating angiosperm species diversity. Hybridization promotes increased genetic and morphological variation, which facilitates speciation, range expansion, and the ecological success of invasive species (Soltis and Soltis, 2009; Cervantes-Díaz et al., 2024). The merging of genomes from distinct origins in first-generation hybrids often induces genomic shock, leading to diverse genetic and epigenetic alterations. When hybrids are fertile generates novel genetic variation that provides raw material for natural selection, enabling the emergence of organisms with varying degrees of adaptation to new environmental conditions. In isolated plant populations harboring genetically unstable hybrid genomes, ongoing natural selection and genetic drift promote the accumulation of genic and chromosomal differences, eventually fostering the development of reproductive isolation mechanisms that enhance genetic divergence and contribute to the formation of new lineages (Rodionov, 2019). Notably, spontaneous hybridization is not uniformly distributed across all plant groups but is more frequent among outcrossing perennials with reproductive strategies that stabilize hybridity, such as agamospermy and vegetative spread (Ellstrand, 1996). Recent genomic and phylogenomic approaches have proven particularly powerful in detecting hybrid origins, introgression, and polyploidization events that are not evident from morphology alone (Moran et al., 2023; Liu et al., 2024).

The recognition and interpretation of hybridization in plant evolution have been greatly advanced by the development of integrative systematics, which combines morphological, cytogenetic, ecological, and molecular data to circumscribe species and evolutionary lineages, particularly in groups affected by reticulate evolution (Schlick-Steiner et al., 2010; Pérez-Ponce de León and Poulin, 2016). In plant groups where hybridization is frequent, reliance on a single line of evidence may obscure evolutionary relationships, as hybrids often exhibit intermediate or mosaic character states and complex patterns of gene flow. By combining multiple lines of evidence, integrative systematics provides a robust framework for identifying hybrid taxa, disentangling reticulate evolutionary histories, and distinguishing between recent hybridization, stabilized hybrid lineages, and incomplete lineage sorting (Solís-Lemus and Ané, 2016; Folk et al., 2018). This approach is particularly critical in plant groups characterized by ongoing gene flow, complex reproductive systems, and rapid diversification, where hybridization and polyploidy play central roles in shaping taxonomic diversity and evolutionary trajectories (Cervantes-Díaz et al., 2024).

### The Nothoscordum system: a lineage prone to genomic and morphological instability

The genus *Nothoscordum* has long been plagued by confusion regarding its circumscription. The large number of described taxa, combined with incomplete species keys and the difficulty of comparing published descriptions with type material and available illustrations, has created a taxonomic impasse. The search for diagnostic morphological characters has been further complicated, making species circumscription particularly challenging (Sassone *et al*., 2025). Reconstruction of molecular phylogenies of the genus has made clear that *N.* sect. *Nothoscordum* has a reticulate evolutionary history, being the most difficult section to circumscribe owing to the wide range of morphological and cytogenetic traits among its species (e.g., Sassone and Giussani, 2018). Our results not only confirm ongoing gene flow between two sympatric species, *N. montevidense* and *N. bonariense* but also provide evidence of backcrossing with the parental species.

While polyploid speciation predominates within *Nothoscordum*, our results suggest that this system represents an early or incipient stage within the continuum of hybrid-driven diversification in perennial plants. This interpretation is supported by the fact that hybrids have not overcome infertility and are likely maintained through asexual propagation. Such a trajectory could potentially lead to homoploid hybrid speciation — defined as the formation of new, reproductively independent lineages derived from hybridization without whole-genome duplication and thus without an increase in ploidy (2*n* = 21 and 2*n* = 25) — as suggested by our GBS results (e.g., Fig. 8). This interpretation is consistent with our initial hypothesis that hybridization within *Nothoscordum* may generate persistent yet evolutionarily dynamic lineages, whose long-term outcomes depend on genomic stabilization, reproductive compatibility, and ecological context rather than on immediate speciation. Evidence from within the genus further supports this view: species such as *N. gramineum* and *N. bivalve* (2*n* = 18), as well as the parental species *N. bonariense* (2*n* = 26) studied here, may represent cases of hybridization without subsequent genome duplication (Souza *et al*., 2019; Palomino *et al*., 1992). Such evolutionary trajectories may have involved extensive chromosomal reorganization following hybridization, including changes in chromosome number through dysploidy or aneuploidization. In these instances, infertility appears to have been overcome, although asexual propagation undoubtedly plays an important role in all such cases.

To fully understand the evolutionary history of these populations, it is necessary to consider the parental species. The shared plastid genome between *N. montevidense* and the hybrid indicates that hybridization occurred with *N. montevidense* as the maternal parent (Fig. 7). This directionality may reflect differences in reproductive compatibility, such as pollen tube growth or ovule development. Previous studies have suggested an allopolyploid origin for *N. bonariense* (Souza et al., 2019; Núñez, 1990), with the current hypothesis proposing that this species combines the two basic chromosome numbers of the genus, *x* = 4 and *x* = 5 (2*n* = 26) (Souza et al., 2019). Based on recent phylogenomic results (Sassone, in prep.), a connection with *N. bivalve* was evident from the plastome phylogeny, suggesting a shared lineage (Fig. 7). The systematics of *N. montevidense* have historically been complicated by its morphological similarity to *N. minarum*; however, these two taxa are now recognized as distinct species based on both base chromosome number and quantitative morphological characters (Guaglianone, 1972; Montes and Nuciari 1987). *Nothoscordum montevidense* has been reported as diploid, tetraploid (as observed here), or octoploid (Montes and Nuciari 1987). Our results further suggest that both parental species share ecological preferences across a large portion of their distribution range (Fig. 4), in contrast to Crosa’s (1972) hypothesis that these two species are predominantly allopatric. Despite significant ecological niche overlap, the hybrid lineage persists alongside the parental lineages, suggesting that ecological differentiation is not a prerequisite for hybrid persistence. Instead, gene flow between the hybrid lineage and the parental species may be constrained by reduced fertility associated with chromosomal imbalance and meiotic irregularities, whereas vegetative propagation may facilitate the persistence of hybrid populations. This supports the view that within *Nothoscordum*, hybrid persistence is driven more by genomic and reproductive dynamics than by niche partitioning.

Moreover, our results suggest that we have captured an evolutionary moment in which the hybrid organisms possess at least two distinct chromosome numbers without yet exhibiting discernible morphological differences. We propose that the hybrid with 2*n* = 21 (1C ≈ 33 pg) backcrossed with *N. bonariense*, generating a karyotype of 2*n* = 25 (1C ≈ 37 pg). The study by Crosa (1972), which we interpret as confirmation of the multiple independent origins of hybridization, identified an individual with 2*n* = 22, also explained as a backcross with *N. bonariense*. This finding points to genomic flexibility within these organisms and the possibility of higher chromosome numbers arising during hybrid formation and backcrossing events. As noted by Núñez (1990), these species are bulbous and perennial, which provides hybrid individuals with sufficient time for natural evolutionary processes to act upon them. It is also worth noting that, despite the wide karyotypic variation documented across the genus (Guerra, 2008), the karyotypes resulting from crosses among the studied plants have not been commonly reported within *Nothoscordum*. This suggests that these chromosomal configurations may represent rare or derived states, potentially reflecting specific evolutionary or genomic dynamics associated with the hybridization process. The meiotic irregularities observed in the hybrid, including lagging chromosomes and micronucleus formation, are consistent with difficulties in chromosome pairing and segregation during meiosis. Such irregularities may result from the combination of parental genomes that differ markedly in chromosome number and karyotype structure (2*n* = 16 in *N. montevidense* and 2*n* = 26 in *N. bonariense*). These differences are expected to challenge homologous chromosome recognition and segregation, potentially contributing to the chromosome-number variation observed among hybrid individuals.

An additional line of evidence supporting ongoing genomic restructuring comes from the analysis of repetitive DNA. Although the abundance of 5S rDNA remained remarkably stable across the parental species and the hybrid, the 35S rDNA fraction showed clear quantitative and qualitative differences. The hybrid exhibited a lower abundance of 35S rDNA than either parental species and a reduced diversity of graph-based rDNA clusters, suggesting differential retention or elimination of parental ribosomal variants following genome merger. Similar patterns have been reported in recently formed hybrids and allopolyploids, where repetitive DNA fractions are among the first genomic components to undergo restructuring through sequence homogenization, copy-number changes, or selective loss of parental variants (Garcia *et al*., 2020). The concordance between repeatome analyses, rDNA-FISH patterns, and the observed meiotic irregularities indicates that the hybrid genome is still undergoing post-hybridization reorganization. Such changes may represent early stages of genome stabilization and contribute to the chromosome-number variation observed among hybrid individuals.

### Evolutionary significance and conservation implications

The current rate of species extinctions, largely driven by anthropogenic activities in the Pampas grasslands, represents a critical threat to biodiversity, particularly for natural populations restricted to small geographic ranges. In this study, we identify a geographically limited area within the genus *Nothoscordum* that exhibits high levels of intraspecific morphological variability despite overall phenotypic similarity among individuals. This pattern suggests the presence of cryptic genetic diversity and highlights the evolutionary significance of these populations as potential reservoirs of unique genetic variation within perennial species, as previously reported for other taxa in the region (e.g. Sassone *et al*., 2021). In the context of increasing habitat loss and fragmentation, our findings underscore the urgent need for targeted conservation strategies aimed at preserving these localized and potentially genetically distinct lineages before irreversible biodiversity loss occurs.

## FUTURE PERSPECTIVES

Future experimental crosses between *N. montevidense* and *N. bonariense* are essential to confirm the hybrid origin of the observed karyotypes, assess the expression of morphological characters, evaluate seed and pollen viability and the stability of hybrid offspring, and conduct detailed analyses of meiotic behavior in these artificial hybrids.

## CONCLUSIONS

Our findings support the hypothesis of low genetic barriers to gene flow within. *Nothoscordum*, providing insight into the evolutionary processes underlying the karyological instability characteristics of this genus. This genetic connectivity offers a compelling explanation for the long-standing taxonomic and systematic challenges within *Nothoscordum*

## DATA AVAILABILITY

Sequence reads for the GBS Illumina runs are stored in the European Nucleotide Archive (ENA, http://www.ebi.ac.uk/ena/data/view/xxx). Sampling locations, morphological matrix and georeferenced localities are provided as supporting information. All other data and scripts are available on xxx (link to repository).

## COLLECTION PERMITS

Botanical collection permits were granted to L.M.G. by the “Ministerio de Gobierno de la provincia de Buenos Aires” under permit “EX 22500-28480-14 DISPO 11; EX-2022-302795470, DISPO-2022-40-GDEBA-DFYFMDAGP”, and its extension “DISPO-2024-38-GDEBA-DFYFMDAGP”; these documents state that “the sovereign rights over natural resources are the exclusive property of the province of Buenos Aires, Argentina. Live specimens were grown in the greenhouse at the Instituto de Botánica Darwinion, and herbarium specimens were deposited there (SI).

## ACKNOWLEDGEMENTS

Alternative version: We gratefully acknowledge Professor Marcelo Guerra for his significant support of this research and his valuable contributions during the early stages of this study.

## AI ACKNOWLEDGEMENTS

ChatAI was used to improve English in the manuscript.

## SUPPLEMENTARY DATA

Supplementary data are available online at https://xxx and consist of the following. Table S1. Voucher information and ENA accessions of the studied taxa. Table S2. Morphological measurements. Table S3. Genomic proportions (%) of repetitive sequence families identified in based on comparative analysis using RepeatExplorer2. Values correspond to the proportion of each repeat family relative to the total genome, obtained from pooled reads representing all analyzed species. Table S4. Results of the Species Distribution Modelling of *Nothoscordum bonariense.* Table S5. Results of the Species Distribution Modelling of *Nothoscordum montevidense*

Figure. S1. Graphical display of the correlation matrix of environmental variables used in the species distribution model for *Nothoscordum montevidense*, using the ‘corrplot’ package in the R enviorenment. Figure. S2. Graphical display of the correlation matrix of environmental variables used in the species distribution model for *Nothoscordum bonariense*, using the ‘corrplot’ package in the R environment. Figure S3. Meiotic analyses revealed irregular meiosis, characterized by the presence of lagging chromosomes during anaphase and the frequent formation of micronuclei. Bar = 5 µm. Figure. S4. Evanno’s plot of the STRUCTURE analysis indicated that K = 2 is the most likely number of genetically distinct populations.

## REFERENCES

Agostinelli C, Lund U. 2025. R package ‘circular’: Circular Statistics (version 0.5-2). https://CRAN.R-project.org/package=circular

Ala N, Bagheri A, Zare H, Himmelbach A, Harpke D, Blattner FR. 2025. Phylogenomics of Western Eurasian *Tilia*: Merging GBS datasets to place the Hyrcanian forest limes. BMC Plant Biology 25: 1370. DOI: 10.1186/s12870-025-07435-4

Andrews S. 2010. FastQC: A Quality Control Tool for High Throughput Sequence Data. http://www.bioinformatics.babraham.ac.uk/projects/fastqc

Bryant D, Moulton V. 2004. Neighbor-Net: An Agglomerative Method for the Construction of Phylogenetic Networks. Molecular Biology and Evolution 21: 255–265.

Bryant D, Moulton V, Spillner A. 2007. Consistency of the neighbor-net algorithm. 640 Algorithms Molecular Biology. 28; 2:8. doi:10.1186/1748-7188-2-8.

Cai L, Xi Z, Amorim AM, Sugumaran M, Rest JS, Liu L, Davis CC. 2019. Widespread ancient whole-genome duplications in Malpighiales coincide with Eocene global climatic upheaval. New Phytologist 221: 565–576. DOI: 10.1111/nph.15357

Cervantes-Díaz L, Mandel JR, Freudenstein JV. 2024. Hybridization, polyploidy and diversification in angiosperms. Botanical Sciences 102: 1–15. DOI: 10.17129/botsci.1420

Crosa O. 1972. Estudios cariológicos en el género Nothoscordum (Liliaceae). Boletin de la Facultad de Agronomía de la Universidad de Montevideo 122: 3–8.

Crosa O. 1974. Un híbrido natural en el género *Nothoscordum* (Liliaceae). Boletín de la Sociedad Argentina de Botánica 15: 471–477.

De Storme N, Mason A. 2014. Plant speciation through chromosome instability and policy change: cellular mechanisms, molecular factors and evolutionary relevance. Current Plant Biology 1: 10–33. DOI: 10.1016/j.cpb.2014.09.002

Doyle JJ, Doyle JL. 1987. A rapid DNA isolation procedure for small quantities of fresh leaf tissue. Phytochemical Bulletin 19: 11–15.

Eaton DAR, Spriggs EL, Park B, Donoghue MJ. 2017. Misconceptions on missing data in RAD-seq phylogenetics with a deep-scale example from flowering plants. Systematic Biology 66: 399–412. DOI: 10.1093/sysbio/syw092

Eaton DAR, Overcast I. 2020. ipyrad: interactive assembly and analysis of RADseq datasets. Bioinformatics 36: 2592–2594. DOI: 10.1093/bioinformatics/btz966

Ellstrand NC, Whitkus R, Rieseberg LH. 1996. Distribution of spontaneous plant hybrids. Proceedings of the National Academy of Sciences of the United States of America 93: 5090–5093. DOI: 10.1073/pnas.93.10.5090

Elshire RJ, Glaubitz JC, Sun Q, et al. 2011. A robust, simple genotyping-by-sequencing (GBS) approach for high diversity species. PLoS ONE 6: e19379. DOI: 10.1371/journal.pone.0019379

Fick SE, Hijmans RJ. 2017. WorldClim 2: new 1-km spatial resolution climate surfaces for global land areas. International Journal of Climatology 37: 4302–4315. DOI: 10.1002/joc.5086

Folk RA, Mandel JR, Freudenstein JV. 2018. Ancestral gene flow and the limits of phylogenetic inference. Systematic Biology 67: 719–734. DOI: 10.1093/sysbio/syy007

Fox J, Weisberg S. 2019. An R Companion to Applied Regression, 3rd edn. Thousand Oaks, CA: Sage. https://www.john-fox.ca/Companion/

Garcia S, Wendel JF, Borowska-Zuchowska N, Aïnouche M, Kuderova A, Kovarik A. 2020. The utility of graph clustering of 5S ribosomal DNA homoeologs in plant allopolyploids, homoploid hybrids, and cryptic introgressants. Frontiers in Plant Science 11: 41. DOI: 10.3389/fpls.2020.00041

Gargiulo R, Kull T, Fay MF. 2021. Effective double-digest RAD sequencing and genotyping despite large genome size. Molecular Ecology Resources 21: 1037–1055. DOI: 10.1111/1755-0998.13314

Gordon A. 2010. Fastx-toolkit. FASTQ/A/short-reads preprocessing tools. https://github.com/agordon/fastx_toolkit

Guaglianone ER. 1972. Sinopsis de las especies de *Ipheion* Raf. y *Nothoscordum* Kunth (Liliáceas) de Entre Rios y regiones vecinas. Darwiniana 17: 159–240.

Guerra M. 2008. Chromosome numbers in plant cytotaxonomy: concepts and implications. Cytogenetic and Genome Research 120: 339–350. DOI: 10.1159/000121083

Hijmans RJ, Etten J v., Cheng J, et al. 2015. ‘raster’: Geographic Data Analysis and Modeling. http://CRAN.R-project.org/package=raster.

Hijmans RJ, Etter J, and others. 2024. dismo: Species Distribution Modeling (version 1.3-16). R package. https://CRAN.R-project.org/package=dismo

Huson DH, Bryant D. 2006. Application of phylogenetic networks in evolutionary studies. Molecular Biology and Evolution 23: 254–267.

Jin JJ, Yu WB, Yang JB, et al. 2020. GetOrganelle: a fast and versatile toolkit for accurate de novo assembly of organelle genomes. Genome Biology 21: 241. DOI: 10.1186/s13059-020-02154-5

Kane N, Sveinsson S, Dempewolf H, Yang JY, Zhang D, Engels JM, Cronk Q. 2012. Ultra-barcoding in cacao (*Theobroma* spp.; Malvaceae) using whole chloroplast genomes and nuclear ribosomal DNA. American Journal of Botany 99: 320–329. DOI: 10.3732/ajb.1100570

Katoh K, Standley DM. 2013. MAFFT multiple sequence alignment software version 7: improvements in performance and usability. Molecular Biology and Evolution 30: 772–780. DOI: 10.1093/molbev/mst010

Lavania UC. 2020. Plant speciation and polyploidy: habitat divergence and environmental perspective. Nucleus 63: 1–5. DOI: 10.1007/s13237-020-00311-6

Liu X, Zhang Y, Wang H, et al. 2024. Phylogenomics reveals extensive hybridization and polyploidy in plants. Annals of Botany 133: mcad089. DOI: 10.1093/aob/mcad089

Moran EV, Soltis PS, Soltis DE. 2023. Genomics of plant speciation. Plant Communications 4: 100530. DOI: 10.1016/j.xplc.2023.100530

Montes L, Nuciari MC. 1987. *Nothoscordum montevidense sensu lato*l: New Polyploid Cytottypes in Argentina. Aliso 11: 635–640.

Nei M, Kumar S. 2000. *Molecular Evolution and Phylogenetics*. Oxford University Press. 352 pp.

Neumann P, Novák P, Hoštáková N, Macas J. 2019. Systematic survey of plant LTR-retrotransposons elucidates phylogenetic relationships of their polyprotein domains and provides a reference for element classification. Mobile DNA 10: 1. DOI: 10.1186/s13100-019-0158-1

Novák P, Neumann P. 2020. Global analysis of repetitive DNA from unassembled sequence reads using RepeatExplorer2. Nature Protocols 15: 3745–3776. DOI: 10.1038/s41596-020-0373-1

Novák P, Neumann P, Pech J, Steinhaisl J, Macas J. 2013. RepeatExplorer: a Galaxy-based web server for genome-wide characterization of eukaryotic repetitive elements from next-generation sequence reads. Bioinformatics 29: 792–793. DOI: 10.1093/bioinformatics/bts048

Núñez O. 1990. Evolución cariotípica en el género *Nothoscordum*. Monografías de la Academia Nacional de Ciencias Exactas, Físicas y Naturales 5: 55–61.

Pellicer J, Hidalgo O, Walker J, Chase MW, Christenhusz MJ, Shackelford G, Fay MF. 2017. Genome size dynamics in tribe Gilliesieae (Amaryllidaceae, subfamily Allioideae) in the context of polyploidy and unusual incidence of Robertsonian translocations. Botanical Journal of the Linnean Society 184: 16–31. DOI: 10.1093/botlinnean/box012

Palomino-Hasbach G, Romo V, Ruenes R. 1992. Centric fissions in metacentric chromosomes of *Nothoscordum bivalve* (Alliaceae) from Mexico. Boletín de la Sociedad Botánica de México 52: 121–124. DOI 10.17129/botsci.1409

Pérez-Ponce de León G, Poulin R. 2016. Taxonomic chauvinism revisited. Systematic Biology 65: 217–226. DOI: 10.1093/sysbio/syv073

Posit team. 2025. RStudio: Integrated Development Environment for R. Boston, MA: Posit Software, PBC. http://www.posit.co/

Pritchard JK, Stephens M, Donnelly P. 2000. Inference of population structure using multilocus genotype data. Genetics 155: 945–959.

R Core Team. 2017. R: A Language and Environment for Statistical Computing. Vienna: R Foundation for Statistical Computing. https://www.R-project.org/

Rieseberg LH, Willis JH. 2007. Plant speciation. Science 317: 910–914. DOI: 10.1126/science.113772

Rodionov AV, Amosova AV, Belyakov EA, et al. 2019. Genetic consequences of interspecific hybridization, its role in speciation and phenotypic diversity of plants. Russian Journal of Genetics 55: 278–294. DOI: 10.1134/S1022795419030141

Sassone AB, Giussani LM, Guerra MR. 2013. Multivariate studies of *Ipheion* (Amaryllidaceae, Allioideae) and related genera. Plant Systematics and Evolution 299: 1561–1575. DOI: 10.1007/s00606-013-0819-5

Sassone AB, López A, Hojsgaard DH, Giussani LM. 2018. A novel indicator of karyotype evolution in the tribe Leucocoryneae (Allioideae, Amaryllidaceae). Journal of Plant Research 131: 211–223.

Sassone AB, Hojsgaard DH, Giussani LM, Brassac J, Blattner FR. 2021. Genomic, karyological and morphological changes of South American garlics (*Ipheion*) provide insights into mechanisms of speciation in the Pampean region. Molecular Ecology 30: 3716–3729. DOI: 10.1111/mec.16009

Sassone AB, Arroyo-Leuenberger S, Moroni P. 2025. Synopsis of the new section *Nothoscordum* sect. *Gracilia* (Amaryllidaceae). Annals of the Missouri Botanical Garden 110: 50–68. DOI: 10.3417/2024023

Schlick-Steiner BC, Steiner FM, Stur E, et al. 2010. Integrative taxonomy: a multisource approach. Trends in Ecology & Evolution 25: 353–361. DOI: 10.1016/j.tree.2010.03.004

Schwarzacher T, Heslop-Harrison P. 2000. *Practical in situ hybridization*. Oxford: BIOS Scientific Publishers.

Solís-Lemus C, Ané C. 2016. Inferring phylogenetic networks with hybridization. PLoS Genetics 12: e1005896. DOI: 10.1371/journal.pgen.1005896

Soltis PS, Soltis DE. 2009. The role of hybridization in plant speciation. Annual Review of Plant Biology 60: 561–588. DOI: 10.1146/annurev.arplant.043008.092039

Soltis PS, Marchant DB, Van de Peer Y, Soltis DE. 2015. Polyploidy and genome evolution in plants. Current Opinion in Genetics & Development 35: 119–125. DOI: 10.1016/j.gde.2015.11.003

Souza G, Crosa O, Speranza P, Guerra M. 2016. Phylogenetic relations in tribe Leucocoryneae (Amaryllidaceae, Allioideae) and the validation of Zoellnerallium based on DNA sequences and cytomolecular data. Botanical Journal of the Linnean Society 182: 811–824. DOI: 10.1111/boj.12463

Souza G, Marques A, Ribeiro T, Dantas L, Speranza P, Guerra M, Crosa O. 2019. Allopolyploidy and extensive rDNA site variation underlie rapid karyotype evolution in *Nothoscordum* section *Nothoscordum* (Amaryllidaceae). Botanical Journal of the Linnean Society. DOI: 10.1093/botlinnean/boz008

Stamatakis A, Ludwig T, Meier H. 2005. RAxML-III: a fast program for maximum likelihood-based inference of large phylogenetic trees. Bioinformatics 21: 456–463. DOI: 10.1093/bioinformatics/bti011

Straub SC, Parks M, Weitemier K, Fishbein M, Cronn RC, Liston A. 2012. Navigating the tip of the genomic iceberg: Next-generation sequencing for plant systematics. American Journal of Botany 99: 349–364. DOI: 10.3732/ajb.1100335

Urbanek S. 2015. rJava: low-level R to Java interface. R package version 0.9–7. http://CRAN.R-project.org/package=rJava

Waminal NE, Pellerin RJ, Kim NS, Jayakodi M, Park JY, Yang TJ, Kim HH. 2018. Rapid and efficient FISH using pre-labeled oligomer probes. Scientific Reports 8: 8224. DOI: 10.1038/s41598-018-26518-8

Wei T, Simko V. 2024. R package ‘corrplot’: visualization of a correlation matrix (version 0.95). https://github.com/taiyun/corrplot

