## Supporting material for "Genomic instability within a sympatric complex of South American garlics (Nothoscordum spp., Amaryllidaceae)"

SUPPLEMENTARY DATA

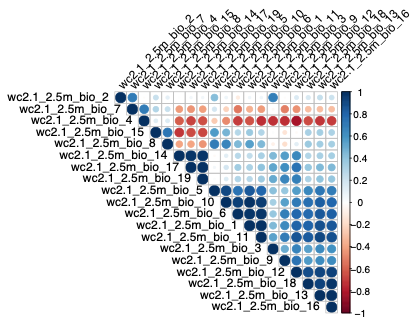
Figure. S1**.** Graphical display of the correlation matrix of environmental variables used in the species distribution model for *Nothoscordum montevidense*, using the 'corrplot' package in the R enviorenment.

| BIO1 | Annual Mean Temperature |
| --- | --- |
| BIO2 | Mean Diurnal Range |
| BIO3 | Isothermality |
| BIO4 | Temperature Seasonality |
| BIO5 | Max Temperature of Warmest Month |
| BIO6 | Min Temperature of Coldest Month |
| BIO7 | Temperature Annual Range |
| BIO8 | Mean Temperature of Wettest Quarter |
| BIO9 | Mean Temperature of Driest Quarter |
| BIO10 | Mean Temperature of Warmest Quarter |
| BIO11 | Mean Temperature of Coldest Quarter |
| BIO12 | Annual Precipitation |
| BIO13 | Precipitation of Wettest Month |
| BIO14 | Precipitation of Driest Month |
| BIO15 | Precipitation Seasonality |
| BIO16 | Precipitation of Wettest Quarter |
| BIO17 | Precipitation of Driest Quarter |
| BIO18 | Precipitation of Warmest Quarter |
| BIO19 | Precipitation of Coldest Quarter |

Figure. S2. Graphical display of the correlation matrix of environmental variables used in the species distribution model for *Nothoscordum bonariense*, using the 'corrplot' package in the R environment.

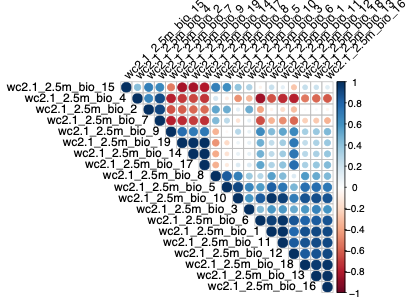

| BIO1 | Annual Mean Temperature |
| --- | --- |
| BIO2 | Mean Diurnal Range |
| BIO3 | Isothermality |
| BIO4 | Temperature Seasonality |
| BIO5 | Max Temperature of Warmest Month |
| BIO6 | Min Temperature of Coldest Month |
| BIO7 | Temperature Annual Range |
| BIO8 | Mean Temperature of Wettest Quarter |
| BIO9 | Mean Temperature of Driest Quarter |
| BIO10 | Mean Temperature of Warmest Quarter |
| BIO11 | Mean Temperature of Coldest Quarter |
| BIO12 | Annual Precipitation |
| BIO13 | Precipitation of Wettest Month |
| BIO14 | Precipitation of Driest Month |
| BIO15 | Precipitation Seasonality |
| BIO16 | Precipitation of Wettest Quarter |
| BIO17 | Precipitation of Driest Quarter |
| BIO18 | Precipitation of Warmest Quarter |
| BIO19 | Precipitation of Coldest Quarter |

Figure S3. Meiotic abnormalities observed in the putative hybrid. **(a)** Irregular metaphase with abnormal chromosome arrangement. **(b, g, j, k)** Lagging chromosomes during anaphase/telophase. **(c, d, f, h, i)** Chromatin bridges connecting segregating chromosome groups. **(e)** Irregular chromosome segregation during telophase. **(l)** Dyad with micronucleus derived from lagging chromosome(s). Scale bar = 5 μm.
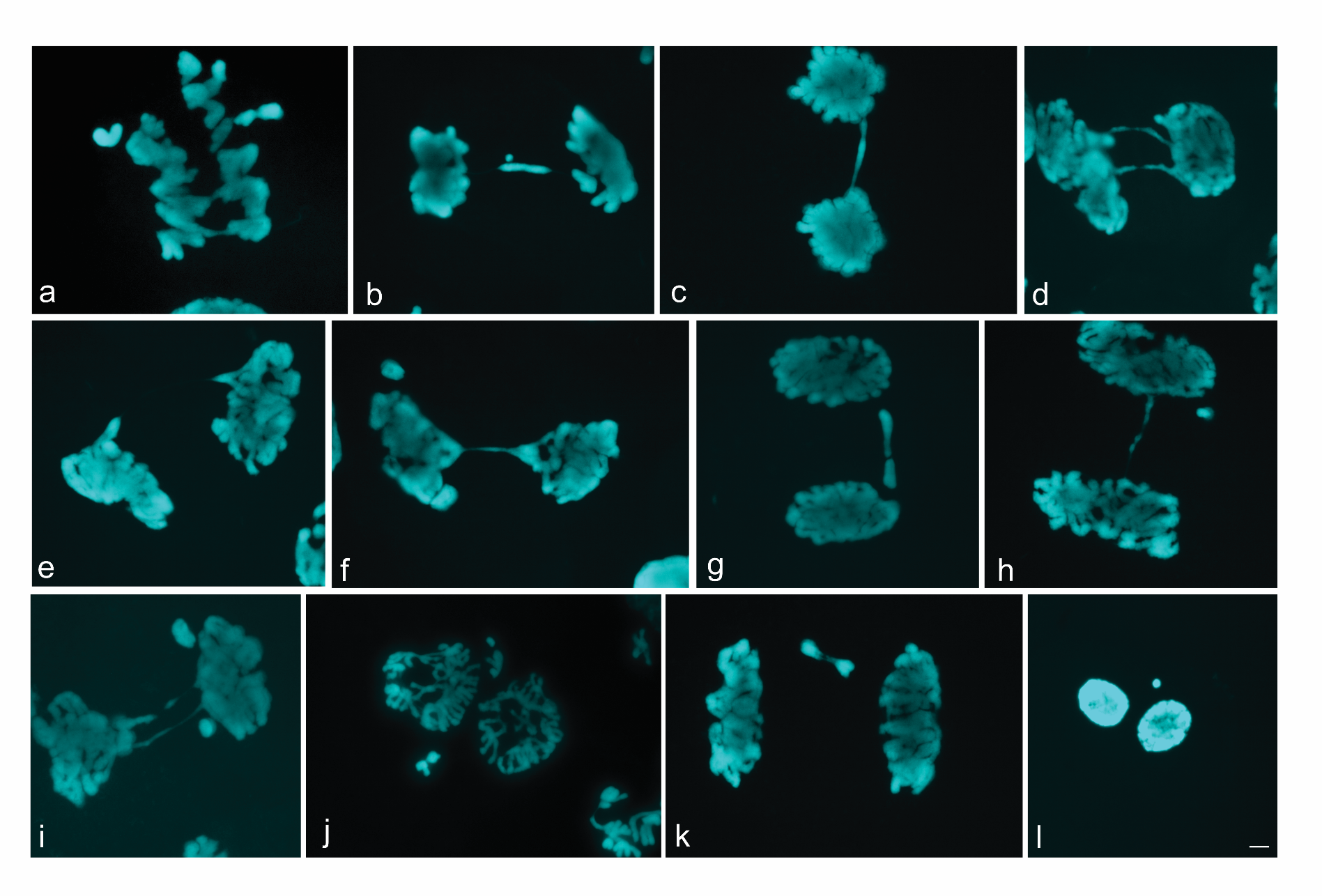

Figure. S4. Evanno's plot of the STRUCTURE analysis indicated that K = 2 is the most likely number of genetically distinct populations.

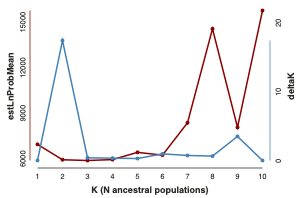

Table S1. Voucher information and ENA accessions of the studied taxa

| **Taxa** | **Origin** | **Voucher** | **Genebank no. Whole chloroplast** | **Genebank no. GBS sequencing** |
| --- | --- | --- | --- | --- |
| *Allium triquetrum* L. | Argentina. Pcia. Bs. As. Tandil (cultivated) | Morrone, O. 6252 (SI) | xxxxxxx | - |
| *Solaria atropurpurea* (Phil.) Ravenna | Argentina. Pcia. Neuquén. Minas | Zuloaga, F. 12510 (SI) | xxxxxxx | - |
| *Miersia chilensis* Lindl*.* | Chile. Biobío. Ñuble | Sassone, A. 47 (SI) | xxxxxxx | - |
| *Latace andina (*Poepp.) Sassone | Chile | Giussani, L. 625 (SI) | xxxxxxx | - |
| *Nothoscordum arenarium Herter* | Uruguay. Soriano | Morrone, O., 6301 (SI) | xxxxxxx | - |
| [*Nothoscordum gracile* (Aiton) Stearn](https://www.ipni.org/n/130726-3) | Chile. Region Metropolitana. Santiago | Sassone, 49 (SI) | xxxxxxx | - |
| *Nothoscordum minarum* | Argentina. Pcia. Entre Ríos | Sassone, A. B. 16 (SI) | xxxxxxx | - |
| *Nothoscordum vittatum* (Griseb.) Ravenna | Uruguay. José Ignacio, laguna Escondida | Giussani, L. 705 (SI) | xxxxxxx | - |
| *Nothoscordum montevidense* | Argentina. Pcia. Bs. As. Magdalena | Giussani, L. 449 (SI) | xxxxxxx | - |
| *Nothoscordum minarum* | Argentina. Pcia. Entre Ríos. Colonia Elia | Giussani, L. 427 a (SI) | xxxxxxx | - |
| *Nothoscordum gramineum* | Chile. Biobío. Ñuble | Giussani, L. 643 (SI) | xxxxxxx | - |
| *Nothoscordum gramineum* | Chile. | From Namgung,J.er al. (2021) | MT348455 | - |
| *Putative hybrid* | Argentina. Pcia. Bs. As. Magdalena | Giussani, L. | xxxxxxx | xxxxxx |
| *Nothoscordum montevidense* | Argentina. Pcia. Bs. As. Tandil | Morrone, O. 6247 (SI) | xxxxxxx | - |
| *Nothoscordum sp.* | Argentina. Pcia. Entre Ríos. Colonia Elia | Giussani, L. 427 b (SI) | xxxxxxx | - |
| *Nothoscordum bonariense* | Argentina. Pcia. Bs. As. Magdalena | Giussani, L. 450 a (SI) | xxxxxxx | xxxxxx |
| *Nothoscordum minarum* | Argentina. Pcia. Entre Ríos. Federación | Sassone, A. 104 (SI) | xxxxxxx | xxxxxx |
| *Nothoscordum bonariense* | Uruguay. Campos bajos frente al Co. Arequita, sobre camino y puente sobre el Río Santa Lucía | Morrone, O. 6311 (SI) | xxxxxxx | - |
| *Nothoscordum bonariense* | Argentina. Pcia. Entre Ríos. Uruguay | Giussani, L., 426 (SI) | xxxxxxx | xxxxxx |
| *Nothoscordum bonariense* | #Argentina. Buenos Aires. La Plata | Sassone, A. 102 (SI) | xxxxxxx | xxxxxx |
| *Nothoscordum bivalve* | México. Durango. El Carmen | Noriega, 16 (SI) | xxxxxxx | - |
| *Nothoscordum bonariense* | Uruguay. Maldonado | Morrone, O. 6312 a (SI) | - | xxxxxx |
| *Nothoscordum bonariense* | Uruguay. Lavalleja | Morrone, O. 6329 a (SI) | - | xxxxxx |
| *Nothoscordum montevidense* | Uruguay. De Rocha a el Pan de Azucar | Morrone, O. s.n. (SI) | - | xxxxxx |
| *Nothoscordum bonariense* | Argentina. Pcia. Entre Ríos. Uruguay | Giussani, L. M. 444 | - | xxxxxx |
| *Nothoscordum montevidense* | Argentina. Pcia. Bs. As. Magdalena | Giussani, L. 449 a (SI) | - | xxxxxx |
| *Nothoscordum montevidense* | Argentina. Pcia. Bs. As. Magdalena | Giussani, L. 449 b (SI) | - | xxxxxx |
| *Nothoscordum montevidense* | Argentina. Pcia. Bs. As. Magdalena | Giussani, L. 449 c (SI) | - | xxxxxx |
| *Nothoscordum montevidense* | Uruguay. Maldonado | Morrone, O. 6312 b (SI) | - | xxxxxx |
| *Nothoscordum montevidense* | Uruguay. Maldonado | Morrone, O. 6312 c (SI) | - | xxxxxx |
| *Nothoscordum bonariense* | Argentina. Pcia. Bs. As. Magdalena | Giussani, L. 450 b (SI) | - | xxxxxx |
| *Putative hybrid* | Argentina. Pcia. Bs. As. Magdalena | Giussani, L. 448 a (SI) | - | xxxxxx |
| *Putative hybrid* | Argentina. Pcia. Bs. As. Magdalena | Giussani, L. 448 b(SI) | - | xxxxxx |
| *Putative hybrid* | Argentina. Pcia. Bs. As. Magdalena | Giussani, L. 448 c (SI) | - | xxxxxx |
| *Putative hybrid* | Argentina. Pcia. Bs. As. Magdalena | Giussani, L. 448 d (SI) | - | xxxxxx |
| *Nothoscordum bonariense* | Argentina. Pcia. Bs. As. Magdalena | Giussani, L. 450 a(SI) | - | xxxxxx |
| *Nothoscordum bonariense* | Argentina. Pcia. Bs. As. Magdalena | Giussani, L. 450 b(SI) | - | xxxxxx |
| *Putative hybrid* | Argentina. Pcia. Bs. As. Magdalena | Giussani, L. 448 c (SI) | - | xxxxxx |
| *Putative hybrid* | Argentina. Pcia. Bs. As. Magdalena | Giussani, L. 448 d (SI) | - | xxxxxx |
| *N. sp 2 (N. sect. Gracilia)* | Uruguay. Pan de Azucar | Sassone, A. 101 a (SI) | - | xxxxxx |
| *N. sp 2 (N. sect. Gracilia)* | Uruguay. Pan de Azucar | Sassone, A. 101 b (SI) | - | xxxxxx |
| *N. sp 2 (N. sect. Gracilia)* | Uruguay. Pan de Azucar | Sassone, A. 101 c (SI) | - | xxxxxx |
| *N. sp 2 (N. sect. Gracilia)* | Uruguay. Pan de Azucar | Sassone, A. 101 d (SI) | - | xxxxxx |

Table S2. Morphological measurements

| **Species** | **BLR** | **BUL** | **LGL** | **PBL** | **PDL** | **SCL** | **CLF** | **FPI** | **NCT** | **PRF** | **NVT** | **FTL** | **OTW** | **ITW** | **OTL** | **ITL** | **CFB** | **FLH** | **ANL** | **ANW** | **OVL** | **OVW** | **STL** | **STG** | **OPL** |
| --- | --- | --- | --- | --- | --- | --- | --- | --- | --- | --- | --- | --- | --- | --- | --- | --- | --- | --- | --- | --- | --- | --- | --- | --- | --- |
| *N. bonariense* | 1 | 0 | 1 | 0 | 4 | 31 | 0 | 8 | 1 | 1 | 0 | 0,2 | 0,4 | 0,2 | 1,3 | 1,1 | 0 | 0,6 | 0,2 | 0,1 | 0,3 | 0,2 | 0,5 | 0 | 4 |
| *N. bonariense* | 1 | 0 | 1 | 0 | 4 | 31 | 0 | 8 | 1 | 1 | 0 | 0,2 | 0,4 | 0,2 | 1,3 | 1,1 | 0 | 0,6 | 0,2 | 0,1 | 0,3 | 0,2 | 0,5 | 0 | 4 |
| *N. bonariense* | 1 | 0 | 1 | 0 | 3,5 | 26 | 0 | 8 | 1 | 1 | 0 | 0,1 | 0,3 | 0,3 | 1,1 | 1,0 | 0,0 | 0,5 | 0,1 | 0,1 | 0,2 | 0,2 | 0,5 | 0 | 4 |
| *intermediate* | 1 | 0 | 1 | 0 | 2,5 | 14 | 2 | 5 | 1 | 1 | 2 | 0,1 | 0,4 | 0,5 | 1,1 | 1 | 0 | 0,5 | 0,2 | 0,1 | 0,2 | 0,2 | 0,4 | 0 | 4 |
| *intermediate* | 1 | 0 | 1 | 0 | 2,5 | 14 | 2 | 5 | 1 | 1 | 2 | 0,1 | 0,4 | 0,5 | 1,1 | 1 | 0 | 0,5 | 0,2 | 0,1 | 0,2 | 0,2 | 0,4 | 0 | 4 |
| *N. montevidense* | 0 | 1 | 1 | 0 | 2,3 | 8 | 1 | 2 | 1 | 1 | 1 | 0,1 | 0,3 | 0,3 | 2 | 1,9 | 0 | 0,4 | 0,2 | 0,1 | 0,2 | 0,15 | 0,3 | 0 | 6 |
| *N. montevidense* | 0 | 1 | 1 | 0 | 2,3 | 8 | 1 | 2 | 1 | 1 | 1 | 0,1 | 0,3 | 0,3 | 2 | 1,9 | 0 | 0,4 | 0,2 | 0,1 | 0,2 | 0,15 | 0,3 | 0 | 6 |

BLR:Bulb with lateral rizomes

BUL:Bulbils

LGL:Ligulated leaves

PBL:Pubescent leaves

PDL:Pedicel length

SCL:Scape length

CLF:Color of the Flower

FPI:Flowers per inflorescence

NCT:Nectar

PRF:Perfume

NVT: Number of tepal nerves?

FTL:Floral tube length

OTW:Outer tepal width

ITW:Inner tepal width

OTL:Outer tepal length

ITL:Inner tepal length

CFB:Connate filaments at base

FLH:Filament length (high)

ANL:Anther length

ANW:Anther width

OVL:Ovary length

OVW:Ovary width

STL:Style length

STG:Stigmata

OPL:Ovules per locule

Table S3. Genomic proportions (%) of repetitive sequence families identified in based on comparative analysis using RepeatExplorer2. Values correspond to the proportion of each repeat family relative to the total genome, obtained from pooled reads representing all analyzed species.

| **Comparative analysis** | *N. montevidense* | *N. montevidense* | **Hybrid** | *N. bonariensis* |
| --- | --- | --- | --- | --- |
| **Class I** |  |  |  |  |
| LINE | 0.13 | 0.12 | **0.11** | 0.11 |
| **LTR** | 11.36 | 11.40 | **11.35** | 11.55 |
| **LTR/Ty1_copia** | 0.05 | 0.05 | **0.04** | 0.05 |
| Ale | 0.20 | 0.20 | **0.21** | 0.19 |
| Alesia | 0.02 | 0.02 | **0.02** | 0.02 |
| Angela | 7.32 | 7.19 | **7.18** | 7.38 |
| Bianca | 0.07 | 0.07 | **0.07** | 0.06 |
| SIRE | 6.76 | 6.82 | **6.89** | 6.79 |
| TAR | 1.35 | 1.38 | **1.38** | 1.32 |
| Tork | 0.86 | 0.86 | **0.88** | 0.89 |
| **LTR/Ty3_gypsy** |  |  |  |  |
| Chromovirus | 1.12 | 1.12 | **1.15** | 1.10 |
| Chromovirus/Tekay | 28.09 | 28.15 | **28.43** | 27.67 |
| Non-chromovirus/OTA/Athila | 2.55 | 2.64 | **2.64** | 2.47 |
| Non-chromovirus/OTA/Tat/Ogre | 16.15 | 16.02 | **15.76** | 16.03 |
| Non-chromovirus/OTA/Tat | 0.03 | 0.02 | **0.02** | 0.03 |
| Non-chromovirus/OTA/Tat/Retand | 7.77 | 7.74 | **7.71** | 7.98 |
| **Class_II** |  |  |  |  |
| Subclass_1/TIR/EnSpm_CACTA | 0.43 | 0.45 | **0.45** | 0.46 |
| Subclass_1/TIR/MuDR_Mutator | 1.42 | 1.50 | **1.52** | 1.53 |
| Subclass_2/Helitron | 0.05 | 0.05 | **0.05** | 0.04 |
| **rDNA** |  |  |  |  |
| 35S rDNA | 0.59 | 0.56 | **0.33** | 0.61 |
| 5S rDNA | 0.02 | 0.02 | **0.02** | 0.02 |
| **Satellite** | 1,68 | 1,66 | **1,61** | 1,69 |
| **Unclassified** | 12.01 | 11.95 | **12.18** | 11.99 |
| **TOTAL** | 69.2 | 68.8 | **69.8** | 69.6 |

Supplementary Table 4. Results of the Species Distribution Modelling of *Nothoscordum bonariense*

| Variable | Percent contribution | Permutation importance |
| --- | --- | --- |
| BIO 14 | 49.4 | 16.6 |
| BIO 4 | 27.6 | 33.1 |
| BIO 19 | 15.9 | 33.6 |
| BIO 7 | 3.8 | 7.2 |
| BIO 15 | 3.3 | 9.6 |

—————

Supplementary Table 5. Results of the Species Distribution Modelling of *Nothoscordum montevidense*

| Variable | Percent contribution | Permutation importance |
| --- | --- | --- |
| BIO 14 | 59.5 | 34.7 |
| BIO 4 | 32.7 | 34.2 |
| BIO 19 | 3.2 | 1.1 |
| BIO 17 | 2.4 | 22.1 |
| BIO 15 | 2.1 | 7.9 |
